# Targeting the Oxysterol Receptor GPR183 to Mitigate Fibrogenesis in Idiopathic Pulmonary Fibrosis

**DOI:** 10.64898/2026.08.09.743811

**Authors:** Minh D. Ngo, Cheng X. Foo, Ziying Hong, Hoang P. L. Uong, Yuanhao Yang, Helle Bielefeldt-Ohmann, Sarah Reed, Edita Ritmejeryte, Lucy Burr, Viviana P. Lutzky, Simon H. Apte, Daniel C. Chambers, Mette M. Rosenkilde, Katharina Ronacher

## Abstract

Idiopathic pulmonary fibrosis (IPF) is a progressive and ultimately fatal lung disease with a median survival of 3-5 years after diagnosis. Current antifibrotic therapies slow disease progression, but do not halt or reverse fibrosis, underscoring the need for new therapies. We identified a dysregulated oxysterol-GPR183 axis as a driver of IPF. Oxidized cholesterols were elevated in lungs from IPF patients, with myofibroblasts representing the dominant source of 7α,25-hydroxycholesterol (7α,25-OHC), the endogenous high affinity ligand for the oxysterol-sensing receptor GPR183. IPF patients had increased GPR183 expression in interstitial and monocyte-like macrophages compared to controls. In a bleomycin-induced model of pulmonary fibrosis genetic deletion of GPR183 reduced disease severity characterized by reduced fibrosis, inflammation, and accumulation of macrophages and myofibroblasts. Pharmacological inhibition of GPR183 with the antagonist NIBR189 attenuated fibrosis when administered preventatively from day 1-7 after bleomycin exposure. Notably, therapeutic treatment with the GPR183 antagonist after commencement of fibrosis development at day 10 post-bleomycin also significantly reduced fibrotic pathology, achieving efficacy comparable to the approved antifibrotic nintedanib. However, the GPR183 antagonist was more potent in reducing inflammation and myofibroblast activation compared to nintedanib. Together, these findings identify an oxysterol-GPR183 signaling axis that contributes to pulmonary fibrogenesis and provide a strong preclinical rationale for targeting GPR183 as a novel therapeutic strategy for IPF.

**One Sentence Summary:** Targeting GPR183 reduced lung fibrosis and inflammation in a preclinical model, supporting GPR183 as a promising new therapy.

## INTRODUCTION

Idiopathic pulmonary fibrosis (IPF) is a progressive interstitial lung disease of unknown cause, characterized by a defective lung repair process and excessive collagen deposition, leading to lung function decline. Despite advances in disease management, the median survival rates of IPF patients range between 3-5 years after diagnosis, highlighting the need to identify novel and more effective treatments (*1–3*). Current understanding of disease pathogenesis indicates that the immune system plays an active role in driving fibrogenesis. Lineage tracing studies have demonstrated that circulating monocytes that are recruited to the injured lung, contribute to disease progression rather than tissue-resident alveolar macrophages (*4, 5*). These monocyte-derived macrophages accumulate within the fibrotic niches and sustain fibrosis through the release of profibrotic mediators including transforming growth factor-β (TGF-β), platelet-derived growth factors (PDGF A and B) and matrix-remodeling enzymes (*6, 7*). Recruited monocytes and macrophages are now representing promising therapeutic targets for IPF (*8, 9*). However, the mechanisms that control the recruitment and positioning of these pathogenic macrophages within fibrotic lung tissue remain incompletely understood.

Emerging evidence suggests that additional guidance systems operate within tissues beside the traditional chemokine pathways. One such pathway is mediated by oxidized cholesterols, acting through the G protein-coupled receptor (GPCR) GPR183, also known as EBI2) (*10, 11*). GPR183-expressing immune cells migrate towards the high-affinity GPR183 agonist 7α,25-dihydroxycholesterol (7α,25-OHC), which is generated from cholesterol through the sequential activity of cholesterol 25-hydroxylase (CH25H) and cytochrome P450 family 7 subfamily B member 1 (CYP7B1) (*10, 11*). Like other chemotactic GPCRs, GPR183 is Gi coupled, and the oxysterol-mediated migration is mediated through Gi and balanced by arresting recruitment (*12, 13*).

We previously demonstrated that GPR183 is an important regulator of monocyte/macrophage recruitment within inflamed lung tissue. We showed that bacterial and viral respiratory infections induce the expression of the oxysterol-synthesizing enzymes CH25H and CYP7b1 in the lung, resulting in local production of 25-OHC and 7α,25-OHC (*14, 15*). Single-cell RNA-sequencing (sc-RNA-seq) data from human bronchoalveolar lavage samples revealed that CH25H, CYP7B1 and GPR183 were upregulated in lung macrophages from COVID-19 patients and associated with disease severity. Importantly, genetic deletion or pharmacological antagonism of GPR183 significantly reduced lung macrophage accumulation, inflammation and disease severity in murine models of Influenza A virus and SARS-CoV-2 infection, positioning GPR183 as a promising drug target for treating airway inflammation (*15*). In pulmonary fungal infection macrophage-derived oxysterols have been shown to be important for spatial positioning of GPR183-expressing T helper 2 cells in inflamed lung tissue (*16*). Lung fibroblasts, the principal effector cells that drive fibrotic tissue remodeling, have recently been identified as a non-hematopoietic source of CH25H and CYP7B1 (*17*). However, despite the growing appreciation of the role of GPR183 in immune-cell trafficking and lung inflammation, the involvement of the GPR183 in pulmonary fibrosis has not previously been investigated.

Here, we show that oxysterols are produced in lungs from IPF patients and that GPR183 ligands accumulate in the lungs during fibrosis progression in a bleomycin-induced IPF mouse model. Genetic deletion of GPR183 protected against pulmonary fibrosis, attenuated myofibroblast activation, and reduced immune cell recruitment during fibrogenesis. Moreover, therapeutic inhibition of GPR183 both prevented fibrosis development and reduced disease severity after fibrosis was established, achieving efficacy comparable to nintedanib. Together, these findings establish GPR183 as a key driver of pulmonary fibrosis and highlight its potential as therapeutic target for the treatment of pulmonary fibrotic disease.

## RESULTS

### Oxysterols are elevated in the lungs of patients with IPF

To investigate whether oxysterol production is altered in IPF, we used targeted Liquid Chromatography-Mass Spectrometry (LC-MS) to measure oxysterol concentrations in bronchoalveolar lavage fluid (BALF) from patients with IPF and IPF patients after receiving a lung transplant (**Fig 1A**). We found that concentrations of 25-OHC were significantly higher in BALF from IPF patients compared to healthy controls (**Fig 1B**). Similarly, 25-OHC was elevated in IPF patients after receiving a lung transplant, suggesting that oxysterol pathways remain active following transplantation. Similarly, concentrations of 27-hydroxycholesterol (27-OHC) were significantly elevated in patients with IPF and lung transplant patients compared with healthy controls (**Fig. 1C**). We attempted to quantify 7α,25-dihydroxycholesterol (7α,25-OHC), the downstream metabolite of 25-OHC, however in this cohort the concentrations of this oxysterol were below the limit of detection.

**Figure 1.**
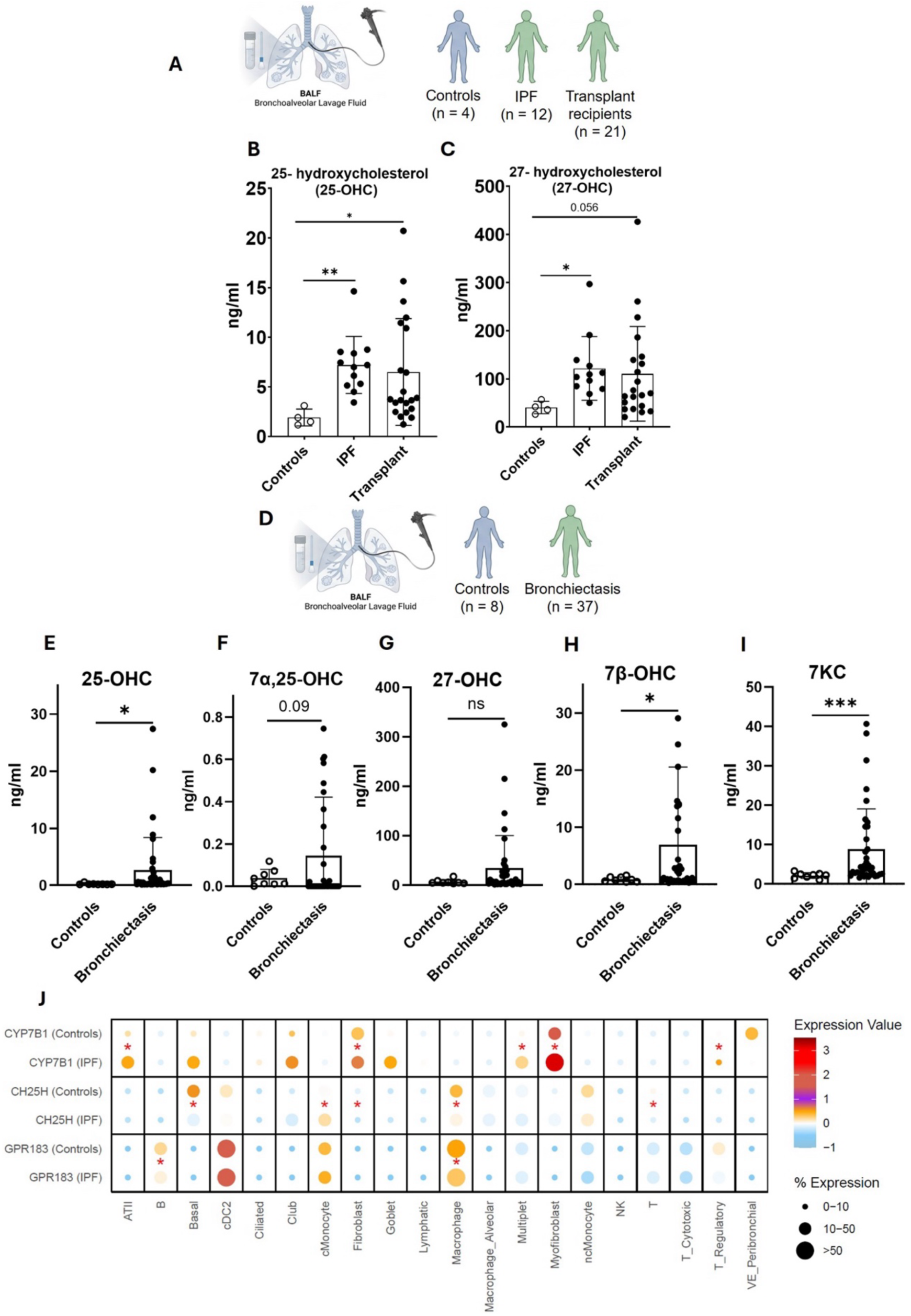
Elevated oxysterols in patients with fibrotic and chronic inflammatory lung diseases. (**A**) Schematic of bronchoalveolar lavage (BALF) collection and patient cohort. BALF samples were obtained from healthy controls (*n* = 4), IPF patients (IPF; *n* = 12), and lung transplant recipients (*n* = 21). Concentrations of (**B**) 25-hydroxycholesterol (25-OHC), and (**C**) 27-dihydroxycholesterol (27-OHC) in BALF were measured by LC-MS. (**D**) Schematic of BALF collection from healthy controls (*n* = 8) and patients with bronchiectasis (BE; *n* = 15). BALF concentrations of (**E**) 25-OHC, (**F**) 7α,25-OHC, (**G**) 27-OHC, (**H**) 7β-OHC, and (**I**) 7 keto-cholesterol (7KC) were measured by LC-MS. Each symbol represents one individual. (**J**) Dot plot showing average normalized expression of CYP7B1, CH25H and GPR183 across major lung cell populations from healthy controls (*n* = 28) and patients with IPF (*n* = 32). Data were derived from a previously published single-cell RNA sequencing dataset (GSE136831)(*18*) comprising 243,472 cells. Cell populations containing ≥1,000 cells were included in the analysis. Dot size indicates the percentage of cells expressing CH25H, CYP7B1 or GPR183 and color denotes average normalized expression. Red asterisks indicate significant differential expression between IPF and control samples. For panels B, C, Kruskal-Wallis analysis followed by Dunn’s multiple-comparison test was used. For panels E-I, statistical significance was assessed using Student’s *t* test or Mann-Whitney *U* test, as appropriate. Data in panel J were analyzed using Wilcoxon rank-sum testing with Bonferroni correction for multiple comparisons. *P < 0.05, **P < 0.01, ***P < 0.001.

To further determine whether oxysterol accumulation is IPF-specific or a broader feature of chronic lung disease and inflammation, we analyzed an independent cohort of patients with bronchiectasis (BE) (**Fig. 1D**). BALF from BE patients compared to control BALF exhibited increased concentrations of 25-OHC and 7α,25-OHC (**Fig. 1E and F**). Patients with BE also exhibited elevated levels of 27-OHC, 7β-hydroxycholesterol (7β-OHC) and 7-ketocholesterol (7KC) compared with healthy controls (**Fig. 1 G-I**). Furthermore, higher concentrations of these oxysterols were associated with poorer lung function, as evidenced by inverse correlations with %FEV1 and %FVC (**Fig. 1A-E**), suggesting that increased oxysterol accumulation is associated with greater disease severity.

To identify cellular compartments associated with the GPR183 ligand 7α,25-OHC in IPF we interrogated an independent scRNA-seq dataset from lung parenchyma samples from 32 IPF patients and 28 healthy controls (*18*). Expression of *CYP7B1*, the enzyme responsible for generating 7α,25-OHC, was detected across several structural and immune cell populations but was most prominently increased in fibroblasts, myofibroblasts and alveolar type II epithelial cells (ATII) from IPF patients compared to controls (**Fig. 1G**). This suggests that 7α,25-OHC is likely increased in patients with IPF relative to healthy controls, however its detection in BALF remains limited by the sensitivity of currently available analytical methods. *CH25H* was increased in fibroblasts and monocytes from IPF patients compared to controls but decreased in macrophages and basal-like cells, while *GPR183* was modestly downregulated in macrophages from IPF patients.

Collectively, these findings demonstrate that oxysterols accumulate within the pulmonary microenvironment in IPF and BE, with particularly myofibroblasts likely contributing to local 7α,25-OHC production.

### The oxysterol-GPR183 pathway is associated with fibrosis development in a mouse model of pulmonary fibrosis

We next investigated the link between oxysterols and lung disease severity in a bleomycin (BLM)-induced murine model of lung fibrosis (**Fig. 2A**).

**Figure 2.**
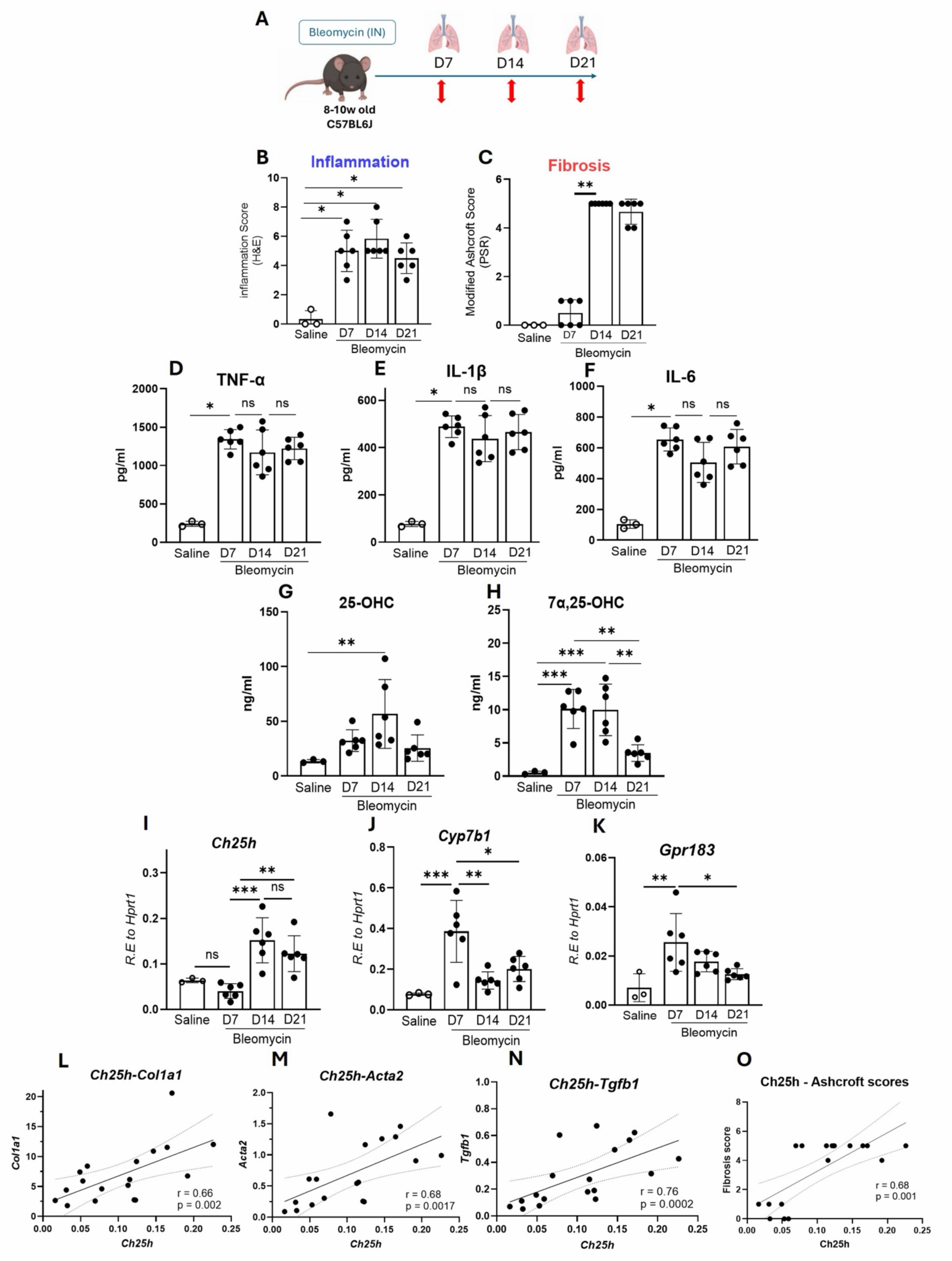
Activation of the oxysterol-GPR183 pathway during bleomycin-induced pulmonary fibrosis. (**A**) Experimental design, 8–10-week-old C57BL/6J mice received a single intranasal (i.n.) dose of bleomycin (BLM, 2.5 U/kg) or saline. Lungs were analyzed at days 7 (D7), 14 (D14), and 21 (D21) post BLM instillations (*n* = 6 per group). Data are from a representative experiment performed independently twice. (**B**) Histological scoring of inflammation on H&E-stained lung sections and (**C**) modified Ashcroft score on Picrosirius Red (PSR)-stained lung sections. (**D-F**) Concentrations of tumor necrosis factor-α (TNF-α), interleukin-1β (IL-1β) and interleukin-6 (IL-6) in lung homogenates measured by ELISA. Quantification of (**G**) 25-OHC and (**H**) 7α,25-OHC following BLM-induced fibrosis in lung homogenates by LC-MS. (**I-K)** Relative lung mRNA expression of Ch25h, Cyp7b1 and Gpr183 by quantitative PCR, normalized to Hprt1 expression. (**L-O**). Correlation analysis between Ch25h expression and the expression of Col1a1, Acta2 and Tgfb1 or Ashcroft scores in BLM-treated lungs from all timepoints (n = 18). Solid lines indicate linear regression, and dotted lines indicate 95% confidence intervals. The Spearman correlation coefficient (*r*) and corresponding *p* values are shown. Data are presented as mean ± SD. Statistical significance was determined by one-way ANOVA with Tukey’s multiple-comparison test. *p< 0.05, **p < 0.01, ns not significance. Col1a1, Collagen type 1 alpha 1 chain; Acta2, Actin alpha 2, smooth muscle; Tgfb1, Transforming growth factor beta 1.

BLM treatment induced lung inflammation by day 7 which persisted up to day 21 as assessed by histopathology (**Fig. 2B**), with fibrosis developing after day 7 as determined by Ashcroft scoring of Picrosirius Red (PSR)-stained lung sections (**Fig. 2C**). Production of the inflammatory cytokines tumor necrosis factor-α (TNF-α), interleukin-1β (IL-1β) and interleukin-6 (IL-6) in the lung was elevated from day 7 to 21 (**Fig. 2D-F**). Concentrations of the oxysterols 25-OHC and 7α,25-OHC were significantly increased in the lungs of BLM-treated mice compared to controls (**Fig. 2G and H**) and associated with the increased mRNA expression of the oxysterol-synthesizing enzymes Ch25h, Cyp7b1 and Gpr183 (**Fig. 2I-K**). As expected, the expression of the fibrogenesis biomarkers Collagen type 1 alpha 1 chain (Col1a1), Fibronectin (Fn1), Transforming growth factor beta (Tgfb1) and Actin alpha 2, smooth muscle (Acta2) were increased in the lung of BLM-treated mice (**Fig. S2**). The expression of Col1a1, Acta2 and Tgfb1 significantly correlated with the mRNA expression of the enzyme Ch25h (**Fig. 2L-N**). Furthermore, Ch25h expression positively correlated with Ashcroft scores indicative of the extent of fibrosis (**Fig. 2O**)

Together, these data demonstrate that GPR183 ligands accumulate in the lungs during fibrosis progression in a BLM-induced IPF mouse model and are associated with markers of lung fibrosis.

### GPR183 deficiency protects mice from pulmonary fibrosis

To investigate the role of GPR183 in the pathogenesis of pulmonary fibrosis, mice deficient in GPR183 (*Gpr183^-/-^*) and age-matched C57BL/6J wild-type mice were administered BLM or saline and disease progression, and severity was assessed at days 7 and 21 post-treatment (**Fig. 3A**). Following BLM administration, C57BL/6J mice exhibited progressive body weight loss (**Fig. 3B**) and increased lung weights (**Fig. 3C**), whereas this response was significantly attenuated in *Gpr183^-/-^* mice, indicating markedly reduced disease severity. Consistent with this, histopathological analysis revealed a reduction in lung inflammation in *Gpr183^-^*^/-^ mice compared with controls (**Fig. 3D and E**) and a significant reduction in the extent of fibrosis (**Fig. 3F and G**). To determine whether the lack of GPR183 influences inflammatory cytokine production during disease development, lung cytokine concentrations were measured. *Gpr183^-/-^* mice exhibited reduced production of IL-1β and IL-6 in the lung (**Fig. 3H and I**), at day 7 following BLM administration. However, cytokine concentrations were comparable between genotypes at later timepoints. Similarly, a trend towards reduced IL-10 and TGF-β1 production was observed in *Gpr183^-/-^* mice at day 7, however this difference did not reach statistical significance (**Fig. 3J and I**).

**Figure 3.**
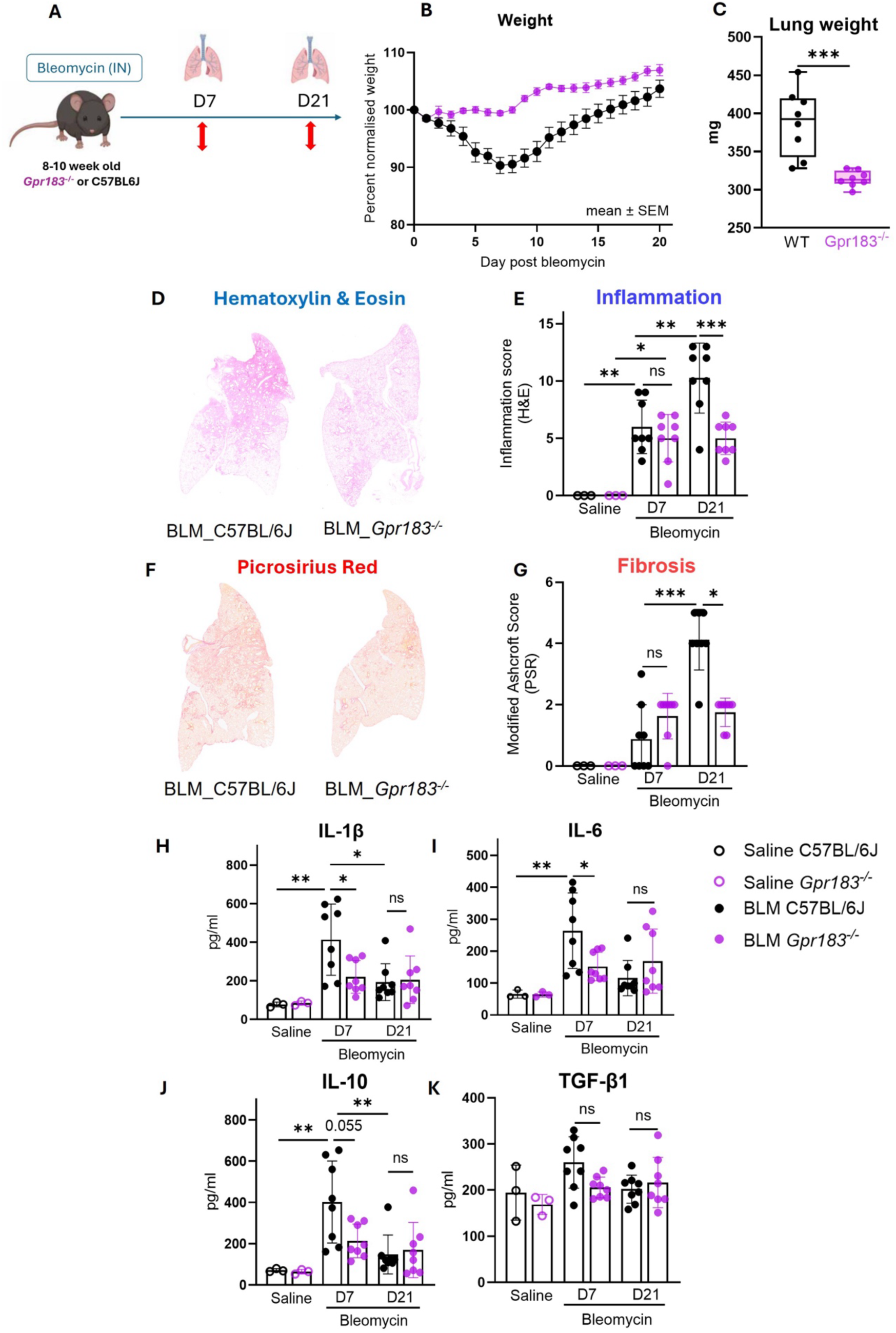
GPR183 deficiency attenuates inflammation and fibrosis. (**A**) Experimental design. *Gpr183*^-/-^ and C57BL/6J mice received a single i.n. administration of BLM (2.5U/kg) and were euthanized at days 7 (D7) or 21 (D21) following treatment (*n* = 8 per genotype per timepoint). Saline-treated mice served as controls (*n* = 3 per genotype). Data are from a representative experiment performed independently three times. (**B**) Body weight changes of C57BL/6J and *Gpr183^-/-^*mice following BLM administration, expressed as percentage of initial weight. (**C**) lung weight measured at day 21 post BLM administration. (**D**) Representative H&E-stained lung sections from C57BL/6J and *Gpr183^−/−^*mice 21 days after bleomycin administration and (**E**) Quantification of inflammation on H&E-stained lung sections. (**F**) Representative PSR-stained lung sections from C57BL/6J and *Gpr183^−/−^* mice 21 days after BLM administration. (**G**) Fibrosis severity quantified using the modified Ashcroft score on PSR-stained lung sections. (**H-K**) Concentrations of IL-1β, IL-6, IL-10 and TGF-β1 in whole lung homogenates at indicated time points, measured by ELISA. Weight graph presented as mean ± SEM. Bar graph data are presented as mean ± SD. Statistical significance in panel C was assessed using an unpaired two-tailed Student’s *t* test. Panel E was analyzed using ordinary one-way ANOVA. Panels G–K were analyzed using Kruskal-Wallis tests followed by Dunn’s multiple-comparison tests. Exact statistical comparisons are indicated in the figure. *p < 0.05, **p < 0.01, ***p < 0.001, ****p < 0.0001; ns, not significant.

These findings indicate that lack of GPR183 protects against BLM-induced inflammation and fibrogenesis in mice.

### GPR183 deficiency attenuates macrophage accumulation and myofibroblast activation in the fibrotic lung

Recruitment of monocytes and macrophages is a key feature of pulmonary fibrosis and has been implicated in disease progression (*7*). Furthermore, we have previously shown that genetic deletion or pharmacological inhibition of GPR183 reduces macrophage accumulation in the lung during respiratory infection with *Mycobacterium tuberculosis* (*14*), Influenza A virus and SARS-CoV-2 (*15*).

To investigate whether this pathway contributes to fibrotic lung disease, lung sections from BLM-treated *Gpr183^-/-^* and wild-type C57BL/6 mice were immunolabelled for the macrophage marker Ionized Calcium-Binding Adapter Molecule 1 (IBA1). IBA1 immunohistochemistry revealed a significantly reduced IBA1^+^ signal in *Gpr183^-/-^* mice compared to control animals at days 7 and 21 post-BLM indicative of reduced macrophage accumulation **(Fig. 4A and B).** We further stained lung sections with an antibody against α-SMA to identify activated myofibroblasts and found significantly fewer α-SMA expressing myofibroblasts in *Gpr183^-/-^* mice **(Fig. 4 C and D)**. IBA1⁺ macrophage accumulation positively correlated with myofibroblast activation marker α-SMA **(Fig. 4E)** and IBA1⁺ macrophages spatially colocalized with α-SMA⁺ myofibroblasts within fibrotic foci **(Fig. 4F)**, suggesting that macrophages contribute to, or sustain, myofibroblast activation.

**Figure 4.**
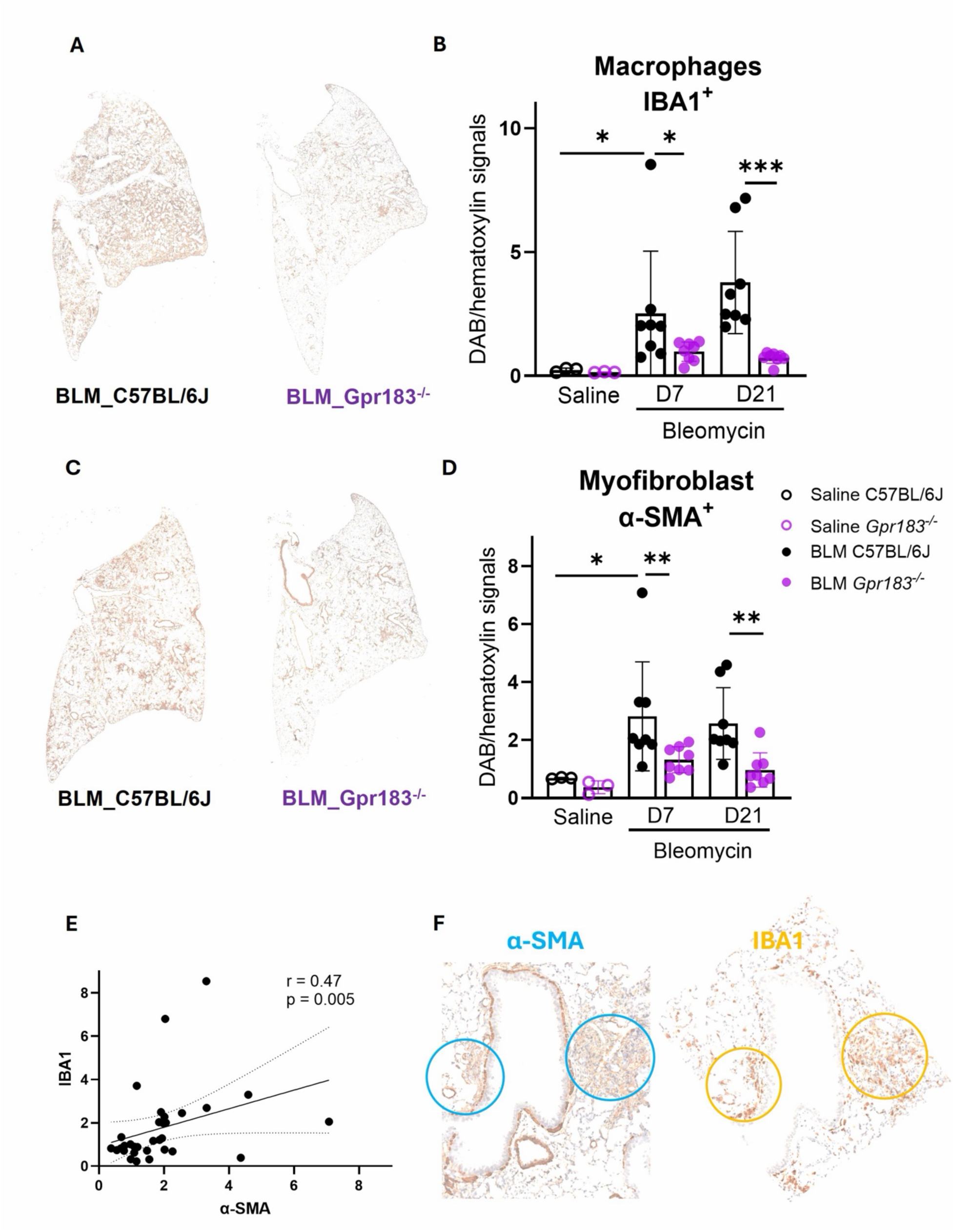
GPR183 deficiency reduces macrophage infiltration and myofibroblast accumulation in the lung after BLM treatment. (**A**) Representative whole-lung immunohistochemical (IHC) staining for IBA1 in lungs from C57BL/6J and *Gpr183^−/−^*mice 21 days after BLM administration (**B**) Quantification of IBA1^+^ staining in lung sections collected from C57BL/6J and *Gpr183^−/−^* mice at days 7 and 21 following BLM administration. (**C**) Representative IHC images of α-SMA in lung tissues at day 21. (**D**) Quantification of α-SMA-positive staining in lung sections collected from WT and *Gpr183^−/−^*mice at days 7 (D7) and 21 (D21) following bleomycin administration. (**E**) Correlation between α-SMA^+^ and IBA1^+^ staining across all BLM-treated mice (n=32). (r = 0.47, *p* = 0.005). (**F**) Representative images showing proximity localization of α-SMA⁺ myofibroblasts (blue circles) and IBA1⁺ macrophages (yellow circles) within fibrotic regions. Statistical significance in panels B and D was assessed using Kruskal-Wallis tests followed by Dunn’s multiple-comparison tests. Correlation analysis in panel E was performed using Spearman’s rank correlation test. Data are presented as mean ± SD. BLM-treated groups (*n* = 8 per genotype at each time point), saline-treated groups (*n* = 3 mice per genotype). Data are from a representative experiment performed independently three times. *p < 0.05, **p < 0.01, ***p < 0.001; ns, not significant. IBA1, ionized calcium-binding adaptor molecule 1. α-SMA, α-smooth muscle actin.

Together, these results demonstrate that loss of GPR183 attenuates macrophage accumulation and diminishes myofibroblast activation in BLM-induced pulmonary fibrosis.

### GPR183 deficiency reduces innate and adaptive immune cell frequencies in the lung

To further investigate how GPR183 influences immune cell dynamics during fibrosis, we analyzed lung immune cell populations by flow cytometry using a gating strategy described in **Fig. S3**. In C57BL/6 mice BLM induced a marked accumulation of interstitial macrophages, monocytes, neutrophils, and T lymphocytes, particularly at day 7 post-BLM challenge, consistent with an acute inflammatory response. In *Gpr183^-/-^* mice the number of interstitial macrophages was significantly reduced compared to C57BL/6 mice at both day 7 and day 21 following BLM treatment (**Fig. 5A**), confirming impaired macrophage accumulation in the absence of GPR183. Similarly, monocyte recruitment was significantly decreased in *Gpr183^-/-^*mice at day 7, although no significant difference was observed at day 21 (**Fig. 5B).** Alveolar macrophage numbers were comparable between genotypes at day 7 but were significantly lower in *Gpr183^-/-^* mice at day 21 (**Fig. 5C**). A reduction in neutrophil infiltration was also observed in *Gpr183^-/-^*mice at day 7, whereas neutrophil numbers were similar between genotypes by day 21 (**Fig. 5D**).

**Figure 5.**
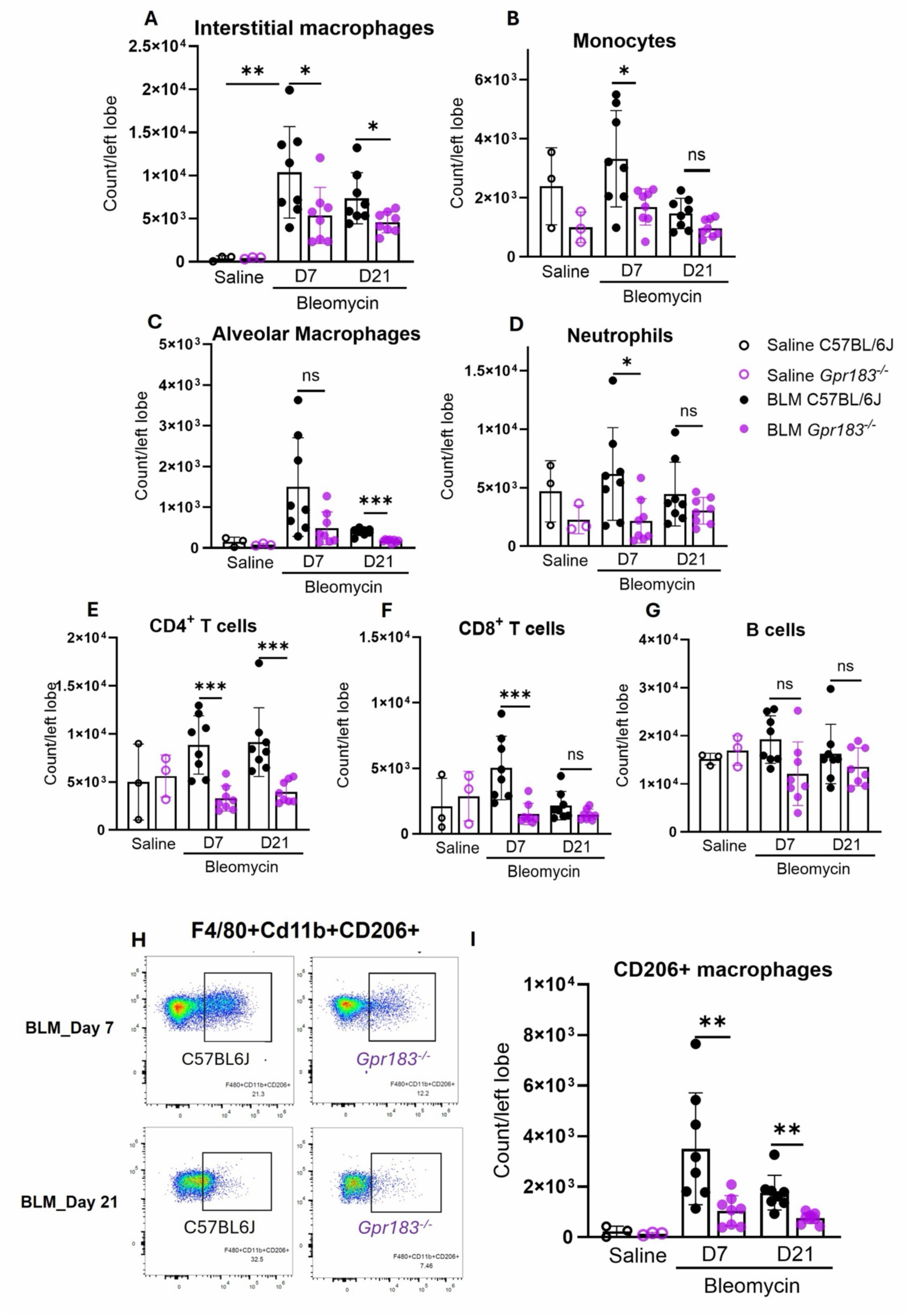
GPR183 deficiency alters myeloid cell populations and macrophage polarization during BLM-induced lung fibrosis. (**A-G**) Quantification of the immune cell populations including interstitial macrophages, monocytes, alveolar macrophages, neutrophils, CD4^+^ T cells, CD8^+^ T cells and B cells in the lung of C57BL6J and *Gpr183*^−/−^mice at day 7 (D7) and day 21 (D21) following BLM administration. (**H**) Representative flow cytometry plots showing F4/80+CD11b+CD206+ macrophage populations in lungs from BLM-treated C57BL/6J and *Gpr183*^-/-^ mice at Day 7 and day 21 post BLM administration. (**I**) Quantification of lung CD206+ macrophages in C57BL/6J and *Gpr183^-/-^* mice at D7 and D21 after BLM administration. Data are from a representative experiment performed independently three times and are presented as mean ± SD. BLM-treated groups (*n* = 8 per genotype at each time point), saline-treated groups (*n* = 3 mice per genotype). *p < 0.05, **p < 0.01, ***p < 0.001; ns, not significant; Student’s *t*-test.

Analysis of lymphocyte populations revealed a marked effect of GPR183 deficiency on T-cell accumulation. Both CD4^+^ and CD8^+^ T cells were significantly reduced in *Gpr183^-/-^* mice at day 7 after BLM treatment (**Fig. 5E and F**). The reduction in CD4^+^ T cells persisted at day 21, whereas CD8^+^ T-cell numbers were no longer significantly different between genotypes at this time point. In contrast, B-cell numbers were not significantly altered by GPR183 deficiency at either time point examined (**Fig. 5G**). BLM administration induced a substantial expansion of CD206^+^ macrophages (CD206^+^F4/80^+^CD11b^+^), a cell population associated with fibrosis progression (*19*), in C57BL/6 mice at both day7 and day21. However, the number of CD206^+^ macrophages was significantly reduced in *Gpr183*^-/-^ mice compared to controls at both time points (**Fig. 5H and I**).

Collectively, these results suggest that lack of GPR183 reduces the immune cell migration and accumulation in the lung and limits the expansion of CD206^+^ macrophage population following lung injury caused by BLM treatment.

### GPR183 antagonism attenuates the development of BLM-induced pulmonary fibrosis

To determine whether pharmacological inhibition of GPR183 can prevent the development of pulmonary fibrosis, mice were treated with the GPR183 antagonist NIBR189 (*20*) or vehicle daily from 24 hours following BLM administration to day 7. Animals were assessed for fibrosis development at the experimental endpoint on day 14 post-BLM treatment **(Fig. 6A)**. Intriguingly, NIBR189-treated mice were protected from BLM-induced body weight loss and exhibited significantly improved recovery compared with vehicle-treated animals **(Fig. 6B)**. At the experimental endpoint, NIBR189-treated mice displayed significantly lower lung weights, consistent with decreased pulmonary inflammation and tissue remodeling (**Fig. 6C**).

**Figure 6.**
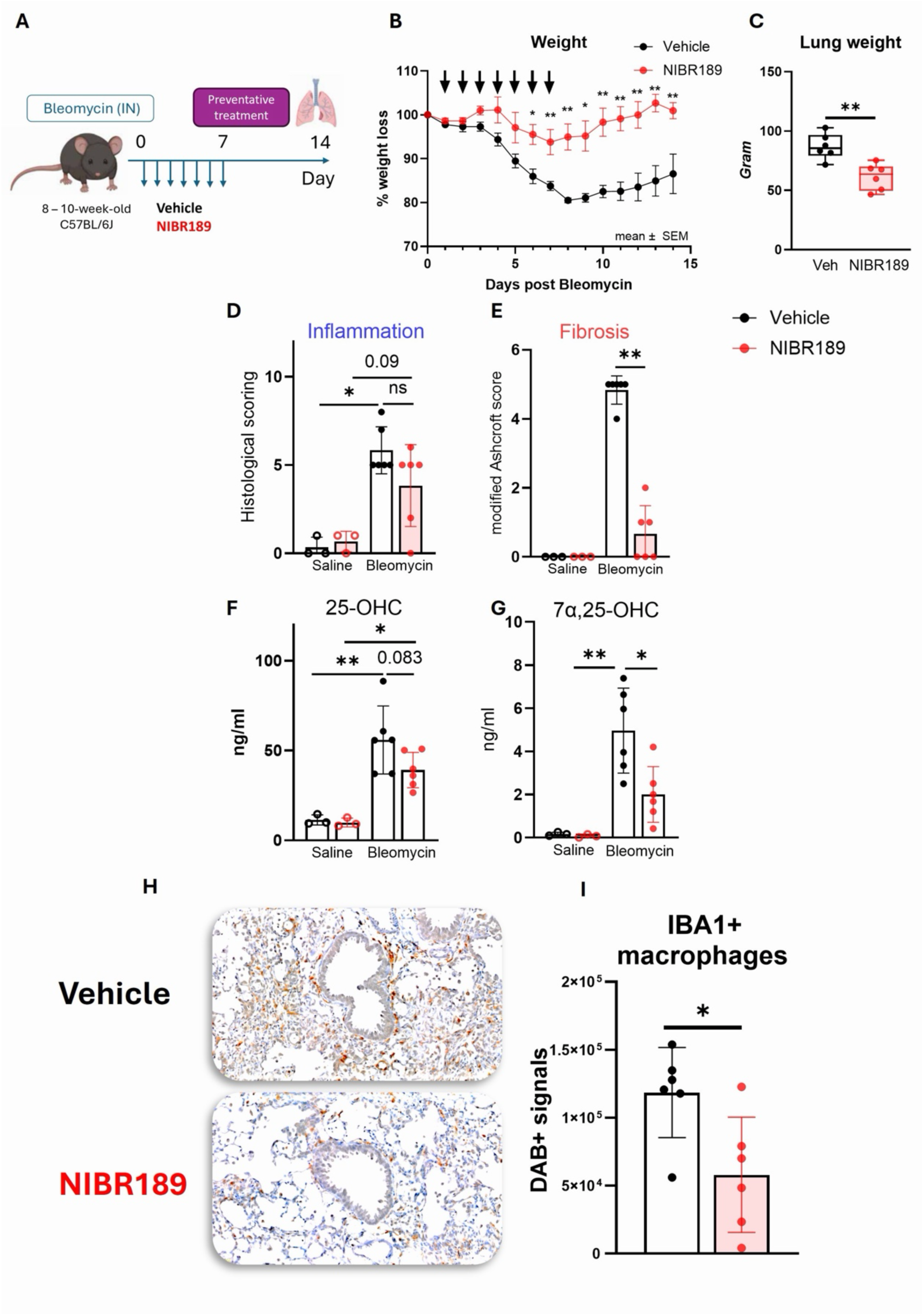
Preventative pharmacological inhibition of GPR183 attenuates BLM-induced pulmonary fibrosis. (**A**) Experimental design, 8-10-week-old C57BL/6J mice received a single i.n. administration of BLM and were treated with the GPR183 antagonist NIBR189 or vehicle daily for the first 7 days post BLM (*n* = 6 per BLM treatment group, saline controls *n* = 3 per group). Mice were euthanized at day 14 post-BLM administration. (**B**) Percentage body weight change following BLM administration in vehicle- and NIBR189-treated mice. (**C**) Lung weights at the experimental endpoint. (**D**) Histopathological inflammation scores determined from H&E-stained lung sections. (**E**) Fibrosis severity quantified using the modified Ashcroft score on Picrosirius Red-stained lung sections. (**F**) Lung concentrations of 25-OHC and (**G**) 7α,25-OHC measured by LC-MS. (**H**) Representative immunohistochemical staining for IBA1 in lungs of vehicle- and NIBR189-treated mice at day 14 following BLM administration. **(I)** Quantification of IBA1 staining expressed as DAB-positive signal intensity. Each symbol represents an individual mouse. Data shown are from a single experiment. Bars indicate mean ± SD. BLM, n = 6 per treatment group; Saline, n = 3 per treatment group Statistical significance was determined using Mann Whitney U test. *p* < 0.05, p < 0.01; ns, not significant.

Although inflammation scores at day 14 were similar between NIBR189-treated and control animals (**Fig. 6D**), NIBR189 significantly reduced fibrosis, as determined by the Ashcroft scoring (**Fig. 6E**). We also observed that NIBR189 treatment reduced pulmonary concentrations of the GPR183 ligand 7α,25-OHC and showed a trend towards reduced 25-OHC concentrations (**Fig. 6F and G**). Since genetic deletion of Gpr183 reduced macrophage accumulation during the fibrotic response to BLM, we next assessed whether pharmacological antagonism produced a similar effect. IHC labelling for IBA1 revealed reduced macrophage accumulation in NIBR189-treated mice compared with vehicle-treated controls (**Fig. 6H and I**).

Together, these findings demonstrate that preventative pharmacological inhibition of GPR183 attenuates BLM-induced pulmonary fibrosis and is associated with reduced macrophage accumulation in the lung.

### GPR183 antagonism improves lung inflammation and fibrosis after fibrosis development

Given that prophylactic treatment of fibrosis is not clinically feasible, we next assessed whether therapeutic treatment initiated after the development of fibrosis could provide similar benefits and limit the progression of established lung fibrosis. Mice were given NIBR189 or vehicle treatment orally from day 10 post-BLM instillation, corresponding to the resolution of the acute inflammatory phase and initiation of fibrogenesis, and treatments continued until day 20 post-BLM. Mice treated with the standard of care antifibrotic nintedanib, a tyrosine kinase inhibitor, served as a positive control for antifibrotic efficacy (**Fig. 7A**).

**Figure 7.**
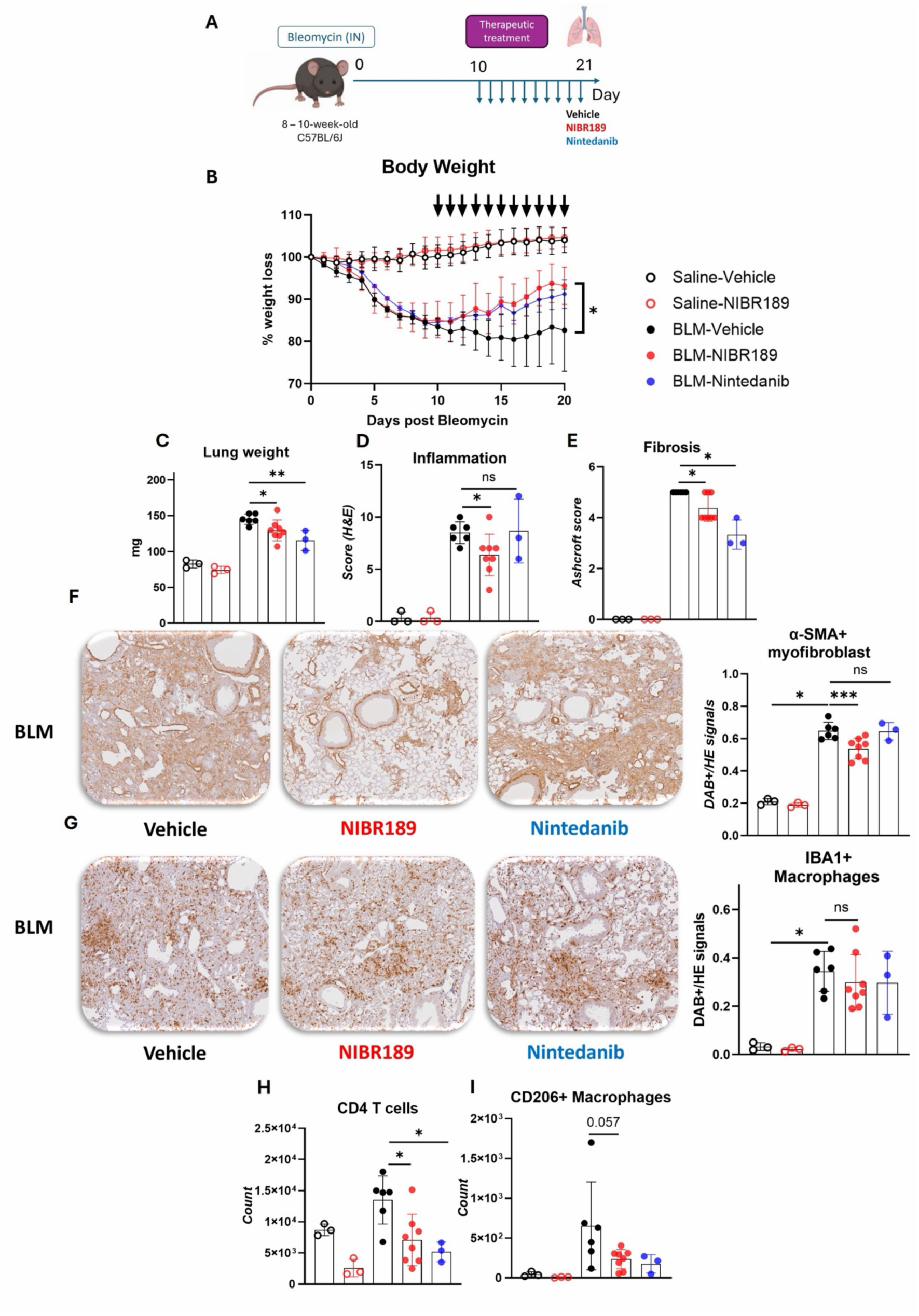
Therapeutic inhibition of GPR183 attenuates established pulmonary fibrosis and modulates immune cell accumulation in the lung. (**A**) Experimental design. Mice received i.n. BLM on day 0. Therapeutic treatment with vehicle (*n* = 6), NIBR189 (*n* = 8), or nintedanib (*n* = 3) commenced on day 10 following BLM administration and continued until day 20. Mice were euthanized on day 21 for endpoint analyses. (**B**) Percentage body weight change following BLM administration in vehicle-, NIBR189-, and nintedanib-treated mice. (**C**) Lung weight at day 21 following BLM administration. (**D**) Histopathological inflammation scores determined from H&E-stained lung sections. (**E**) Fibrosis severity quantified using the modified Ashcroft score on Picrosirius Red-stained lung sections. (**F**) Representative immunohistochemical staining and quantification of α-smooth muscle actin (α-SMA) in lung sections collected at day 21 following BLM administration. (**G**) Representative immunohistochemical staining and quantification of ionized calcium-binding adaptor molecule 1 (IBA1) in lung sections collected at day 21 following BLM administration. (**H**) Frequencies of pulmonary CD4⁺ T cells and (**I**) CD206⁺ macrophages determined by flow cytometry. Bars indicate mean ± SD. Statistical significance was determined using two-way ANOVA with Sidak’s multiple-comparison test for body weight measurements and Kruskal-Wallis analysis followed by Dunn’s multiple-comparison test for endpoint analyses. Exact statistical comparisons are shown in the figure. *p < 0.05, **p < 0.01, \*\*\***p** < 0.001; ns, not significant.

We found that NIBR189-treated mice exhibited improved recovery of body weight compared with vehicle-treated controls. Body weight recovery in NIBR189-treated mice was comparable to that observed in the nintedanib treatment group (**Fig. 7B**). Consistent with these findings, lung weights were significantly reduced at day 21 in NIBR189-treated mice compared with vehicle-treated control mice (**Fig. 7C**). Therapeutic administration of NIBR189 significantly reduced inflammation scores, while inflammation scores were similar between vehicle and nintedanib-treated animals (**Fig. 7D**). Assessment of collagen deposition by PSR staining further demonstrated a substantial reduction in fibrotic lesions in NIBR189-treated mice compared with vehicle-treated mice, producing an antifibrotic effect similar to that achieved with nintedanib (**Fig. 7E**). Histologically, NIBR189 treatment significantly reduced the α-SMA signal, indicating decreased myofibroblast accumulation and activation (**Fig. 7F**). IHC labelling for IBA1 did not reach significant differences in macrophage numbers between treatment groups (**Fig. 7G**).

To investigate whether therapeutic GPR183 inhibition influenced pulmonary immune cell populations, lung leukocytes were analyzed by flow cytometry. We found that NIBR189 treatment reduced the accumulation of CD4⁺ T cells with a trend toward fewer CD206⁺ macrophages compared with vehicle-treated controls (**Fig. 7H and I**), partially recapitulating the immunological phenotype observed in *Gpr183^-/-^* mice. However, no significant difference was found in other immune cell subsets (**Fig. S4**). Despite the reduction in inflammation scores, endpoint concentrations of inflammatory cytokines, including TGF-β1, were not significantly altered by NIBR189 treatment (**Fig. S5**). A trend towards reduced *Col1a1* expression was observed in animals treated with NIBR189. Expression of the fibrosis-associated genes *Fn1, Acta2*, and *Tgfb1* not changed by NIBR189 treatment, while nintedanib significantly increased *Tgfb1* expression (**Fig. S6**).

Collectively, these findings demonstrate that therapeutic inhibition of GPR183 after the acute inflammatory phase is sufficient to reduce myofibroblast expansion, lung inflammation, and fibrosis in murine BLM-induced fibrosis, suggesting that GPR183 represents a promising therapeutic target for the treatment of established pulmonary fibrosis (**Fig. 8**).

**Figure 8.**
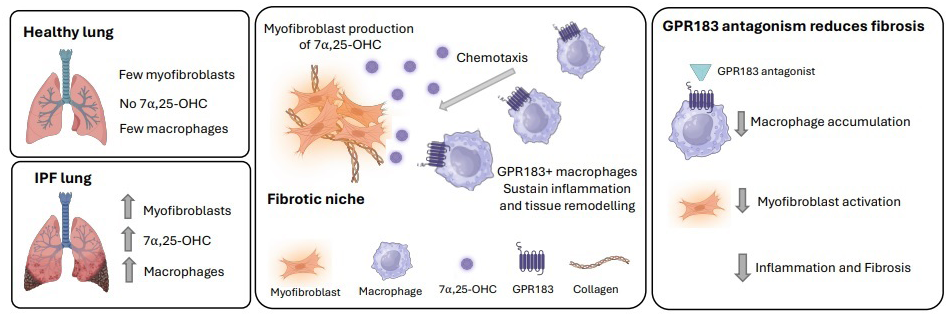
Schematic overview of the main findings. During IPF myofibroblasts produce the high affinity endogenous ligand for GPR183 7α,25-OHC which chemotactically attracts peripheral monocytes/macrophages to the fibrotic niche. GPR183^+^ macrophages sustain inflammation and tissue remodeling. GPR183 antagonism reduces macrophage accumulation, myofibroblast activation, inflammation and fibrogenesis.

## DISCUSSION

The recruitment of circulating monocytes and macrophages into the fibrotic niches is recognized as central event in IPF pathogenesis. Pro-fibrotic macrophages are increasingly viewed as attractive therapeutic targets in IPF (*4, 6, 9*). Furthermore, single-cell and spatial transcriptomic studies have highlighted the importance of microenvironmental niches in sustaining fibrosis in which fibroblasts, epithelial cells, and immune cells engage in coordinated interactions (*9, 21*). Our study provides evidence that the oxysterol-GPR183 axis represents a critical spatial signaling mechanism linking fibroblasts and infiltrating monocytes/macrophages within these niches.

We found that oxysterol production is elevated in the diseased human lung in both IPF and BE, providing important evidence that enhanced oxysterol synthesis is a common feature of chronic lung injury. Although we detected the high-affinity GPR183 ligand 7α,25-OHC in clinical samples from BE patients, concentrations were very low and fell below the limit of detection in the IPF cohort, representing a limitation of this study. One possible explanation is that slow oxidative degradation occurred during long-term storage of bio banked samples, even at -80°C. Nevertheless, the sc-RNA Seq data showed increased expression of CYP7B1 mRNA in fibroblasts, myofibroblasts and ATII cells and provides indirect evidence that 7α,25-OHC production is elevated in IPF lungs. Consistent with this interpretation, our animal studies demonstrate that increased Cyp7b1 expression in the lung is associated with increased 7α,25-OHC. While we did not directly measure CYP7B1 enzymatic activity, these findings collectively support enhanced 7α,25-OHC biosynthesis in IPF.

Consistent with our findings in human IPF, we observed activation of the oxysterol-GPR183 signaling axis during the development of fibrosis in the BLM-induced mouse model. Genetic loss-of-function studies provided direct evidence that GPR183 plays a functional role in disease pathogenesis. Compared with wild-type controls, *Gpr183^-/-^* mice exhibited reduced weight loss, lower inflammatory cytokine production, diminished immune cell infiltration, and markedly attenuated fibrosis. Together, these findings indicate that GPR183 contributes to both the inflammatory and fibrotic phases of disease and suggest that oxysterol-driven GPR183 signaling is an important regulator of pulmonary fibrogenesis.

Mechanistically, our findings identify macrophages as key cellular targets of the GPR183 pathway in pulmonary fibrosis. Histological analysis revealed reduced numbers of IBA1-positive macrophages and α-SMA-positive myofibroblasts in *Gpr183^-/-^* lungs. In fibrotic wild-type lungs, these populations were spatially associated and significantly correlated, supporting a relationship between macrophage accumulation and fibroblast activation. Given the established role of macrophages in promoting fibrosis through secretion of TGF-β1 and other profibrotic mediators, as well as through direct interactions with fibroblasts, our data suggest that oxysterol-GPR183 signaling contributes to the maintenance of a profibrotic macrophage-fibroblast niche. We propose that local production of 7α,25-OHC within the injured lung facilitates the recruitment and development of GPR183-expressing tissue-resident macrophages (*17*), which subsequently drive fibroblast activation, myofibroblast differentiation, and extracellular matrix deposition, thereby reinforcing fibrotic tissue remodeling.

Flow cytometric analyses provided further insight into the immune cell populations regulated by GPR183 in the lung. The absence of major differences in immune cell composition between C56BL/6 and *Gpr183^-/-^*mice at baseline suggests that GPR183 is largely dispensable for pulmonary immune homeostasis but becomes functionally important during tissue injury and inflammation. Following lung injury, loss of GPR183 selectively affected specific immune cell subsets. The most consistent changes were observed in interstitial macrophages, which were reduced at both early and late stages of disease, while monocytes were reduced during the early inflammatory phase. These findings are consistent with our previous studies in infection models demonstrating that GPR183 is required for the rapid recruitment of monocytes and macrophages to the lung (*14, 15*). Others have similarly reported an essential role for GPR183 in eosinophil trafficking (*22*). In contrast, GPR183 appears to be redundant for T-cell positioning within the lung under infectious conditions (*15, 16, 23*). It was therefore unexpected that *Gpr183^-/-^*exhibited reduced numbers of both CD4^+^ and CD8^+^ T cells in the fibrosis model, a finding that was not observed in our infection models (*15*). This observation suggests that the mechanisms governing lymphocyte accumulation during chronic fibrotic remodeling may differ from those operating during acute infection and may involve indirect effects on cellular recruitment, retention or survival within the injured lung. Likewise, the reduction in neutrophils observed in *Gpr183^-/-^*mice is likely to occur through indirect mechanisms, as neutrophils do not express GPR183. Together, these data position monocytes and macrophages as primary cellular targets of GPR183 signaling in the fibrotic lung, with downstream consequences for the recruitment and/or maintenance of additional cell populations that contribute to disease progression.

Notably, pharmacological inhibition of GPR183 reduced disease severity in both preventive and therapeutic treatment settings. When compared with the current standard-of-care antifibrotic agent nintedanib, GPR183 antagonism demonstrated comparable efficacy in reducing fibrosis, while providing superior suppression of pulmonary inflammation and myofibroblast accumulation. These findings suggest that targeting GPR183 may offer advantages over existing antifibrotic therapies by intervening earlier in the pathogenic cascade. Whereas nintedanib primarily targets fibroblasts, myofibroblasts and endothelial cells through inhibition of multiple receptor tyrosine kinases, including PDGFR, FGR and VEGF, GPR183 antagonism acts further upstream by limiting the recruitment and positioning of profibrotic monocytes and macrophages within the injured lung. Given the restricted expression of GPR183 to specific immune cell populations, therapeutic inhibition may result in a more selective mechanism of action and potentially a more favorable safety profile than broad-spectrum tyrosine kinase inhibition. Although this remains to be demonstrated experimentally, the limited cellular distribution of GPR183 raises the possibility of a wider therapeutic window with fewer off-target effects. From a translational perspective, GPR183 is an especially attractive target because it belongs to the GPCR superfamily, one of the most successful classes of drug targets in modern medicine (*24*). The extensive clinical experience with GPCR-directed therapies highlights the tractability of this receptor family and supports the feasibility of developing GPR183-targeted therapeutics for pulmonary fibrosis. Importantly, despite their distinct mechanisms of action, GPR183 antagonists and nintedanib may not be redundant. Instead, targeting both the upstream immune drivers of fibrosis and the downstream fibroproliferative response could provide complementary therapeutic benefits, making combination therapy an attractive avenue for future investigation.

In conclusion, our study identifies the oxysterol-GPR183 pathway as a previously unrecognized contributor to IPF pathogenesis. The findings of the study establish a mechanistic link between oxysterol signaling and fibrotic remodeling and provide strong preclinical evidence that targeting GPR183 represents a promising therapeutic strategy for IP. GPR183 inhibition may offer a novel approach to disrupt the immune-fibrotic circuitry that drives progressive lung fibrosis.

## MATERIALS AND METHODS

### Study Design

The objective of this study was to investigate the role of oxysterols and the oxysterol-sensing receptor GPR183 in lung fibrosis in both human studies and animal models. Human bronchoalveolar lavage fluid (BALF) samples were obtained from consenting participants of case control cohorts according to international guidelines as previously described (*25*). Archived frozen BALF samples from patients with IPF (*n* = 12), IPF patients post lung transplantation (n=21) and healthy control subjects (*n* = 4) were collected and stored at The Prince Charles Hospital (Brisbane, Australia). Sample collection and use were approved by the Metro North Hospital and Health Service Human Research Ethics Committee (HREC/2023/MNHB/94455). A second cohort of archived BALF samples from patients with bronchiectasis (BE, *n* = 34) and healthy control subjects (*n* = 12) was obtained from the Mater Biobank (Mater Adult Hospital, Brisbane, Australia) under approval from the Mater Misericordiae Limited Human Research Ethics Committee (EXMT/MML/119748). All studies were conducted in accordance with relevant institutional guidelines and national regulations.

All animal studies were conducted in compliance with protocols approved by the animal ethics committee of the University of Queensland (2023/AE000628). Mice were randomly assigned to groups. Experimental endpoints were predetermined and are indicated in the text and figure legends. A single operator veterinary pathologist assessed lung sections and was blinded to the treatment allocation and animal genotypes. Study participant numbers and numbers of animals can be found in then figure legends. The number of repeat experiments is indicated in the figure legends. No outliers were excluded.

### Bioinformatic analysis of human single-cell RNA sequencing data

A previously published scRNA-seq data set from lung parenchyma samples obtained from 32 IPF patients and 28 healthy controls (*18*) (GEO accession number GSE136831) was analyzed. Differential expression of *GPR183*, *CH25H* and *CYP7B1* was assessed within individual cell-type clusters using the Wilcoxon rank-sum test implemented in the *Seurat* (v5.2.0) R package (*26*), comparing IPF with healthy controls in the lung parenchyma dataset. Cell-type clusters were included in the analysis if they contained at least 1,000 cells to maintain sufficient statistical power. Genes were considered for testing if they exhibited an absolute average log2 fold change >0.25 and were expressed in at least 5% of cells in either comparison group. Statistical significance was determined after Bonferroni correction for multiple testing.

### Mouse model of pulmonary fibrosis

C57BL/6J mice were purchased from the Jackson Laboratory. *Gpr183^-/-^*(Gpr183^tm1LEX^) where obtained from Lexicon Pharmaceuticals and back-crossed to a C57BL/6J background. All mice were maintained in house and used for experiments at 8–10-weeks of age.

To induce lung fibrosis, mice were anesthetized by intraperitoneal injection of ketamine (120 mg/kg) and xylazine (20 mg/ kg) and received intranasal (IN) instillation of a single dose of bleomycin sulfate (BLM) (1.5-2.5U/kg, MP Biomedicals, #0219030610) dissolved in 50μL sterile saline as previously described (*27*). Dose of BLM was optimized for each lot to achieve consistent fibrotic response and comparable body-weight loss across experiments. Control mice received an equivalent volume of sterile saline. Mice were weighed daily to monitor their health status, and animals that lost more than 15% of their initial body weight were euthanized according to ethical endpoints. At the experimental endpoint (day 7, 14 or 21), mice were euthanized by carbon dioxide asphyxiation, and lungs were collected for downstream analyses.

The GPR183 antagonist NIBR189 (7.6 mg/kg; Merck, #SML1981) or vehicle (0.5% carboxymethylcellulose (CMC) and 0.05% Tween-80) was administered via oral gavage twice daily. Nintedanib (40 mg/kg; Merck, #SML2848), an approved anti-fibrotic therapy for IPF, was administered once daily by oral gavage in the same vehicle formulation. Mice were euthanized at indicated timepoints and lungs were harvested for histopathology, immunohistochemistry, flow cytometry, RNA analysis and ELISA.

As male mice exhibit a more robust fibrotic response to bleomycin-induced lung injury compared to female (*28, 29*) males were used for all experiments except the preventative NIBR189 study, which was performed in female mice.

### Liquid Chromatography-Mass Spectrometry (LC-MS)

Liquid extraction method: Oxysterols were extracted from BALF samples using a modified methanol–chloroform extraction protocol. Briefly, methanol (100%, LC-MS grade) was added sample (1:1, v:v ratio) and vortexed for 30 seconds. Then, chloroform was added to achieve a PBS:methanol:chloroform ratio of 1:1:2. After vortexing and centrifugation (3,500 rpm, 5 min), the organic phase was collected. Chloroform extraction was repeated twice, and pooled organic fractions were dried under nitrogen, reconstituted in 50 µL methanol, and analyzed by mass spectrometry.

Solid extraction Method: Oxysterol extraction from lung tissue samples was performed as previously published (*14, 15*), by using dichloromethane:methanol (1:1) solution containing 50 µg/mL butylated hydroxytoluene (BHT). Samples were flushed with N₂, sonicated (30°C, 10 min), and centrifuged (3,500 rpm, 5 min). Following liquid–liquid extraction of the supernatant with DPBS (3,500 rpm, 5 min at 25 °C), the organic phase was collected and dried under N₂. Oxysterols were purified by aminopropyl solid-phase extraction (SPE), eluted with chloroform:methanol (23:1), dried under N₂, and resuspended in 100 µL of 90% methanol containing 0.1% DMSO prior to analysis and placed in an ultrasonic bath for 5 min at 30°C.

### Mass spectrometry analysis

Calibration curves (0.39–800 ng/mL) were prepared using authentic standards of 25-OHC, 7α,25-OHC, 27-OHC, 24-OHC, 7β-OHC, and 7-ketocholesterol obtained from Sigma-Aldrich and Avanti. Analyte concentrations were quantified using the corresponding calibration curves. Each analytical batch included calibration standards, QC samples, and unknown samples.

Samples were analyzed on AB Sciex QTRAP® 5500 (AB SCIEX, Redwood City, CA) mass spectrometer coupled to a Shimadzu Nexera 2 UHPLC. Oxysterols were separated on a Kinetex Pentafluorophenyl (PFP) column (100 x 2.1mm, 2.6µM, Phenomenex) using water (0.1% formic acid) and acetonitrile (0.1% formic acid) as mobile phases A and B, respectively, at a flow rate of 0.5 mL/min. The gradient was held at 40% B for 1.3 min, increased linearly to 60% B over 15 min and then to 99% B over 1 min. The column was washed at 99% B for 2 min and re-equilibrated at 40% B for 1 min before the next injection. The column and autosampler temperatures were maintained at 50°C and 15°C, respectively.

Oxysterols were detected in positive ESI mode using scheduled MRM on an AB Sciex QTRAP® 5500 mass spectrometer with a Turbo V™ DuoSpray source (550°C, 5500 V). Curtain gas (CUR), ion source gas 1 (GS1), and ion source gas 2 (GS2) were set to 30, 65, and 50, respectively. Data were acquired using a 90 s detection window and 0.5 s cycle time, with Q1 and Q3 operated at unit resolution. Compound-specific MRM transitions and optimized DP, CE, EP, and CXP parameters used for data acquisition are provided in Supplementary Table 1. Data were processed in SCIEX OS v 4.0.0.8559, with automated peak integration and reviewed manually. As 24-OHC, 25-OHC, and 27-OHC shared identical MRM transitions, identification was confirmed by retention time using authentic reference standards.

### Hematoxylin & eosin and Picrosirius red staining of lung sections

The left lung lobe was inflated with 10% neutral-buffered formalin and fixed in the same solution for 24 h at room temperature. Fixed tissues were routine processed for paraffin embedding and preparation of hematoxylin and eosin (H&E) stained sections. For assessment of collagen deposition, sections were stained with Picrosirius Red (PSR) solution. H&E- and PSR-stained sections were imaged by brightfield microscopy and used for histopathological assessment of inflammation and fibrosis.

### Histopathological scoring of lung sections

Histopathological assessment for inflammation and fibrosis was performed by an independent veterinary pathologist blinded to genotype and treatment allocation. Lung inflammation was evaluated on H&E-stained sections by assessing changes within the interstitium, alveoli, and bronchioles. Each parameter was scored using a semi-quantitative scale as follows: 0, no change; 1, minimal change; 2, mild change; 3, moderate change; 4, severe change affecting <50% of the lung lobe; and 5, severe change affecting >50% of the lung lobe. Scores for the individual compartments were summed to generate a total inflammation score. The fibrotic state of murine lungs was evaluated using the modified Ashcroft scoring system as previously described (*30*). Briefly, low and higher power magnification images were obtained from PSR stained and scanned murine lungs. Qualitative evaluation of the images was performed by an independent veterinary pathologist, blinded to the treatment groups, scoring each image from a scale of 0-8 where 0 is normal lung to 8 being most fibrotic.

### Immunohistochemistry staining

After tissue sections were deparaffinized and rehydrated, antigen retrieval was performed in 10 mM sodium citrate buffer (pH 6.0) using a decloaking chamber at 105°C for 20 min, followed by cooling for 20 min at room temperature. Endogenous peroxidase activity was quenched with 3% H₂O₂ in TBS for 15 min, and sections were blocked with 1% BSA in TBS for 15 min at room temperature. Sections were incubated with rabbit anti-Iba1 antibody (Novachem, SO33866; 1:1,000) or rabbit α-SMA (Abcam, AB124964; 1:1000) for 2h at room temperature, followed by an HRP-conjugated anti-rabbit secondary antibody for 30 min. Immunoreactivity was visualized using 3,3′-diaminobenzidine (DAB) with 60 second incubation for IBA1 staining and 20 second for α-SMA staining. Nuclei were counterstained with Gill’s hematoxylin. Subsequently, sections were dehydrated through graded ethanol, cleared in xylene, and mounted with DPX mounting medium. Positive staining was quantified by Qupath analysis software (V0.6.0).

### Flow cytometry

Tissue preparation for flow cytometry analysis was performed as described previously (*15*). Briefly, single-cell suspensions were prepared from mouse lungs by enzymatic digestion with Liberase (50 μL per sample; Roche) in phenol red-free RPMI/DMEM at 37°C with gentle agitation (300 rpm) for 20 min. Digested tissue was passed through a 70-μm cell strainer, mechanically dissociated using a syringe plunger, washed with cold phenol red-free medium, and centrifuged at 400 × g for 5 min at 4°C. Red blood cells were lysed using 1× RBC lysis buffer for 1.5 min. Cells were resuspended in MACS buffer and stained with a fixable viability dye (ViaDye Red, 1:2,000; Biotium) for 15 min on ice, followed by Fc receptor blockade (1:200) for 20 min and incubation with fluorochrome-conjugated antibodies for 30 min on ice in the dark. A complete list of antibodies used for flow cytometric analyses is provided in Supplementary Table 2. Cells were then fixed in 10% neutral buffered formalin, washed, resuspended in MACS buffer and stored at 4°C protected from light until acquisition. Flow cytometry data were acquired on a Cytek aurora flow cytometer, and data were analyzed using FlowJo software (V10, BD Biosciences) Among pulmonary macrophage populations, tissue-resident alveolar macrophages were identified as CD45^+^Ly6G⁻CD11c⁺Siglec-F⁺ cells. Interstitial macrophages were defined as CD45⁺Ly6G⁻F4/80⁺CD11b⁺Siglec-F⁻ cells. CD206⁺ macrophages were identified as CD45^+^F4/80⁺CD11b⁺CD206⁺ cells. Inflammatory monocytes were identified as CD45⁺CD11b⁺Ly6C⁺ cells. Neutrophils were defined as CD45⁺CD11b⁺Ly6G⁺ cells. T cells were identified as CD45⁺CD3⁺ lymphocytes and subsequently classified into CD4⁺ and CD8⁺ subsets. B cells were identified as CD45⁺B220⁺ cells. Representative gating strategies are provided in Supplementary Fig. S1 (*4*).

### RNA extractions and qPCR

RNA was extracted using Trizol (Thermo Fisher Scientific, 10296028) and the ISOLATE II RNA Mini Kit (Meridian Bioscience, BIO-52073). Complementary DNA (cDNA) was synthesized from total RNA using the Tetro cDNA synthesis kit (Meridian Bioscience, BIO-65043) according to the manufacturer’s instructions. Quantitative real-time PCR (qPCR) was performed using SensiFAST^TM^ SYBR Lo-ROX kit (Meridian Bioscience, BIO-94050) and a QuantStudio™ 3 Real-Time PCR System machine (Applied Biosystems™). Primer sequences are listed in Supplementary table 3.

### Statistical analysis

All data are presented as mean ± standard deviation (SD) unless otherwise stated in the figure legends. The same statistical framework applies to both human and mouse studies. Data were first tested for normality using the Shapiro-Wilk test. For comparisons between two groups, either an unpaired two-tailed Student’s *t*-test or Mann-Whitney U test was used as appropriate. Comparisons among three or more groups were performed using one-way ANOVA or Kruskal-Wallis test followed by appropriate multiple-comparison correction depending on data distribution. The statistical tests used for each dataset are indicated in the corresponding figure legends. A *P* value < 0.05 was considered statistically significant. Quantitative data were assembled and analyzed using GraphPad Prism version 10.0 (GraphPad Software, San Diego, CA, USA) The number of biological replicates (depicted as *n*) is indicated in the corresponding figure legends.

## List of Supplementary Materials

Fig. S1 to S6

Table S1 to S3

## Acknowledgments

The authors acknowledge the Translational Research Institute for providing an excellent research environment and core facilities that enabled this research. We also thank the staff of the Biological Resources Facility, Histology Facility, and Flow Cytometry Facility for their technical and administrative support. Additionally, the authors thank the staff and patients who have donated their time and biological samples to the Mater Research Biobank and the Prince Charles Hospital.

## Funding

CREATE Hope Fellowship (MDN)

LINC Grant, Translational Research Institute (MDN, LB)

Prince Charles Hospital Foundation (VL, SHA, DC, KR)

Australian NHMRC 2019167 (KR)

Mater Foundation (KR)

The Translational Research Institute is supported by the Australian Government

## Author contributions

Conceptualization: MDN, DCC, MMR, KR

Methodology: MDN, CXF, ZH, HPLU, YY, HBO, SR, ER, MMR, LB, VPL, SHA, DCC

Investigation: MDN, CXF, ZH, HPLU, YY, HBO, VPL, SHA, DCC, MMR, KR

Visualization: MDN, CXF, YY

Funding acquisition: MDN, LB, VPL, SHA, DCC, KR

Project administration: KR

Supervision: KR

Writing – original draft: MDN, KR

Writing – review & editing: MDN, CXF, ZH, HPLU, YY, HBO, SR, ER, LB, VPL, SHA, DCC, MMR, KR

## Data and materials availability

All data are available in the main text or the supplementary materials.

## References

1. G. Raghu, M. Remy-Jardin, J. L. Myers, L. Richeldi, C. J. Ryerson, D. J. Lederer, J. Behr, V. Cottin, S. K. Danoff, F. Morell, K. R. Flaherty, A. Wells, F. J. Martinez, A. Azuma, T. J. Bice, D. Bouros, K. K. Brown, H. R. Collard, A. Duggal, L. Galvin, Y. Inoue, R. G. Jenkins, T. Johkoh, E. A. Kazerooni, M. Kitaichi, S. L. Knight, G. Mansour, A. G. Nicholson, S. N. J. Pipavath, I. Buendia-Roldan, M. Selman, W. D. Travis, S. Walsh, K. C. Wilson, E. R. S. J. R. S. American Thoracic Society, S. Latin American Thoracic, Diagnosis of Idiopathic Pulmonary Fibrosis. An Official ATS/ERS/JRS/ALAT Clinical Practice Guideline. Am J Respir Crit Care Med 198, e44–e68 (2018).

2. Q. Zheng, I. A. Cox, J. A. Campbell, Q. Xia, P. Otahal, B. de Graaff, T. J. Corte, A. K. Y. Teoh, E. H. Walters, A. J. Palmer, Mortality and survival in idiopathic pulmonary fibrosis: a systematic review and meta-analysis. ERJ Open Res 8, (2022).

3. D. J. Lederer, F. J. Martinez, Idiopathic Pulmonary Fibrosis. N Engl J Med 378, 1811–1823 (2018).

4. A. V. Misharin, L. Morales-Nebreda, P. A. Reyfman, C. M. Cuda, J. M. Walter, A. C. McQuattie-Pimentel, C. I. Chen, K. R. Anekalla, N. Joshi, K. J. N. Williams, H. Abdala-Valencia, T. J. Yacoub, M. Chi, S. Chiu, F. J. Gonzalez-Gonzalez, K. Gates, A. P. Lam, T. T. Nicholson, P. J. Homan, S. Soberanes, S. Dominguez, V. K. Morgan, R. Saber, A. Shaffer, M. Hinchcliff, S. A. Marshall, A. Bharat, S. Berdnikovs, S. M. Bhorade, E. T. Bartom, R. I. Morimoto, W. E. Balch, J. I. Sznajder, N. S. Chandel, G. M. Mutlu, M. Jain, C. J. Gottardi, B. D. Singer, K. M. Ridge, N. Bagheri, A. Shilatifard, G. R. S. Budinger, H. Perlman, Monocyte-derived alveolar macrophages drive lung fibrosis and persist in the lung over the life span. J Exp Med 214, 2387–2404 (2017).

5. H. Aegerter, B. N. Lambrecht, C. V. Jakubzick, Biology of lung macrophages in health and disease. Immunity 55, 1564–1580 (2022).

6. J. I. Bailey, C. H. Puritz, K. J. Senkow, N. S. Markov, E. Diaz, E. Jonasson, Z. Yu, S. Swaminathan, Z. Lu, S. Fenske, R. A. Grant, H. Abdala-Valencia, R. J. Mylvaganam, A. Ludwig, J. Miller, R. I. Cumming, R. M. Tighe, K. M. Gowdy, R. Kalhan, M. Jain, A. Bharat, C. Kurihara, R. San Jose Estepar, R. San Jose Estepar, G. R. Washko, A. Shilatifard, J. I. Sznajder, K. M. Ridge, G. R. S. Budinger, R. Braun, A. V. Misharin, M. A. Sala, Profibrotic monocyte-derived alveolar macrophages are expanded in patients with persistent respiratory symptoms and radiographic abnormalities after COVID-19. Nat Immunol 25, 2097–2109 (2024).

7. S. Wang, J. Li, C. Wu, Z. Lei, T. Wang, X. Huang, S. Zhang, Y. Liu, X. Bi, F. Zheng, X. Zhu, Z. Huang, X. Yi, Single-Cell RNA Sequencing Reveals Monocyte-Derived Interstitial Macrophages with a Pro-Fibrotic Phenotype in Bleomycin-Induced Pulmonary Fibrosis. Int J Mol Sci 25, (2024).

8. H. Cui, S. Banerjee, N. Xie, M. Hussain, A. Jaiswal, H. Liu, T. Kulkarni, V. B. Antony, R. M. Liu, M. Colonna, G. Liu, TREM2 promotes lung fibrosis via controlling alveolar macrophage survival and pro-fibrotic activity. Nat Commun 16, 1761–1776 (2025).

9. N. Joshi, S. Watanabe, R. Verma, R. P. Jablonski, C. I. Chen, P. Cheresh, N. S. Markov, P. A. Reyfman, A. C. McQuattie-Pimentel, L. Sichizya, Z. Lu, R. Piseaux-Aillon, D. Kirchenbuechler, A. S. Flozak, C. J. Gottardi, C. M. Cuda, H. Perlman, M. Jain, D. W. Kamp, G. R. S. Budinger, A. V. Misharin, A spatially restricted fibrotic niche in pulmonary fibrosis is sustained by M-CSF/M-CSFR signalling in monocyte-derived alveolar macrophages. Eur Respir J 55, 1900646–1900666 (2020).

10. S. Hannedouche, J. Zhang, T. Yi, W. Shen, D. Nguyen, J. P. Pereira, D. Guerini, B. U. Baumgarten, S. Roggo, B. Wen, R. Knochenmuss, S. Noel, F. Gessier, L. M. Kelly, M. Vanek, S. Laurent, I. Preuss, C. Miault, I. Christen, R. Karuna, W. Li, D. I. Koo, T. Suply, C. Schmedt, E. C. Peters, R. Falchetto, A. Katopodis, C. Spanka, M. O. Roy, M. Detheux, Y. A. Chen, P. G. Schultz, C. Y. Cho, K. Seuwen, J. G. Cyster, A. W. Sailer, Oxysterols direct immune cell migration via EBI2. Nature 475, 524–527 (2011).

11. C. Liu, X. V. Yang, J. Wu, C. Kuei, N. S. Mani, L. Zhang, J. Yu, S. W. Sutton, N. Qin, H. Banie, L. Karlsson, S. Sun, T. W. Lovenberg, Oxysterols direct B-cell migration through EBI2. Nature 475, 519–523 (2011).

12. M. M. Rosenkilde, T. Benned-Jensen, H. Andersen, P. J. Holst, T. N. Kledal, H. R. Luttichau, J. K. Larsen, J. P. Christensen, T. W. Schwartz, Molecular pharmacological phenotyping of EBI2. An orphan seven-transmembrane receptor with constitutive activity. J Biol Chem 281, 13199–13208 (2006).

13. V. M. S. Kjaer, V. Daugvilaite, T. M. Stepniewski, C. M. Madsen, A. S. Jorgensen, K. R. Bhuskute, A. Inoue, T. Ulven, T. Benned-Jensen, S. A. Hjorth, G. M. Hjorto, E. V. Moo, J. Selent, M. M. Rosenkilde, Migration mediated by the oxysterol receptor GPR183 depends on arrestin coupling but not receptor internalization. Sci Signal 16, eabl4283 (2023).

14. M. D. Ngo, S. Bartlett, H. Bielefeldt-Ohmann, C. X. Foo, R. Sinha, B. J. Arachchige, S. Reed, T. Mandrup-Poulsen, M. M. Rosenkilde, K. Ronacher, A Blunted GPR183/Oxysterol Axis During Dysglycemia Results in Delayed Recruitment of Macrophages to the Lung During Mycobacterium tuberculosis Infection. J Infect Dis 225, 2219–2228 (2022).

15. C. X. Foo, S. Bartlett, K. Y. Chew, M. D. Ngo, H. Bielefeldt-Ohmann, B. J. Arachchige, B. Matthews, S. Reed, R. Wang, C. Smith, M. J. Sweet, L. Burr, K. Bisht, S. Shatunova, J. E. Sinclair, R. Parry, Y. Yang, J. P. Levesque, A. Khromykh, M. M. Rosenkilde, K. R. Short, K. Ronacher, GPR183 antagonism reduces macrophage infiltration in influenza and SARS-CoV-2 infection. Eur Respir J 61, (2023).

16. Y. Zheng, H. E. Dobson, M. Jean Pierre, L. B. LeBlanc, C. U. Onyishi, D. P. Golec, N. Carrillo, A. Deewan, C. A. Rivera, E. Ansaldo, P. L. Schwartzberg, E. V. Dang, Macrophage-derived oxysterols organize T helper 2 cells to suppress type 1 inflammation during pulmonary fungal infection. Sci Immunol 11, eaee0990 (2026).

17. L. Bub, E. Evren, S. Verwaerde, C. Ruscitti, D. Vanneste, P. Ghosh, Y. Gao, N. Sleiers, R. Deng, M. Lopez Montes, K. Howley, R. La Rocca, A. Niehrs, V. Glaros, M. Reina-Campos, B. Dahlen, A. Smed-Sorensen, H. Lund, T. Kreslavsky, N. K. Bjorkstrom, A. Reboldi, A. Bossios, T. Marichal, B. N. Lambrecht, T. Willinger, Sensing of metabolic signals via GPR183 promotes occupation of lung macrophage niches by monocytes. J Exp Med 223, 1–26 (2026).

18. T. S. Adams, J. C. Schupp, S. Poli, E. A. Ayaub, N. Neumark, F. Ahangari, S. G. Chu, B. A. Raby, G. DeIuliis, M. Januszyk, Q. Duan, H. A. Arnett, A. Siddiqui, G. R. Washko, R. Homer, X. Yan, I. O. Rosas, N. Kaminski, Single-cell RNA-seq reveals ectopic and aberrant lung-resident cell populations in idiopathic pulmonary fibrosis. Sci Adv 6, eaba1983 (2020).

19. L. Pommerolle, G. Beltramo, L. Biziorek, M. Truchi, A. M. M. Dias, L. Dondaine, J. Tanguy, N. Pernet, V. Goncalves, A. Bouchard, M. Monterrat, G. Savary, N. Pottier, K. Ask, M. R. J. Kolb, B. Mari, C. Garrido, B. Collin, P. Bonniaud, O. Burgy, F. Goirand, P. S. Bellaye, CD206(+) macrophages are relevant non-invasive imaging biomarkers and therapeutic targets in experimental lung fibrosis. Thorax 79, 1124–1135 (2024).

20. F. Gessier, I. Preuss, H. Yin, M. M. Rosenkilde, S. Laurent, R. Endres, Y. A. Chen, T. H. Marsilje, K. Seuwen, D. G. Nguyen, A. W. Sailer, Identification and characterization of small molecule modulators of the Epstein-Barr virus-induced gene 2 (EBI2) receptor. J Med Chem 57, 3358–3368 (2014).

21. A. C. Habermann, A. J. Gutierrez, L. T. Bui, S. L. Yahn, N. I. Winters, C. L. Calvi, L. Peter, M. I. Chung, C. J. Taylor, C. Jetter, L. Raju, J. Roberson, G. Ding, L. Wood, J. M. S. Sucre, B. W. Richmond, A. P. Serezani, W. J. McDonnell, S. B. Mallal, M. J. Bacchetta, J. E. Loyd, C. M. Shaver, L. B. Ware, R. Bremner, R. Walia, T. S. Blackwell, N. E. Banovich, J. A. Kropski, Single-cell RNA sequencing reveals profibrotic roles of distinct epithelial and mesenchymal lineages in pulmonary fibrosis. Sci Adv 6, 1972–1987 (2020).

22. A. C. Bohrer, E. Castro, C. E. Tocheny, M. Assmann, B. Schwarz, E. Bohrnsen, M. A. Makiya, F. Legrand, K. L. Hilligan, P. J. Baker, F. Torres-Juarez, Z. Hu, H. Ma, L. Wang, L. Niu, Z. Wen, S. H. Lee, O. Kamenyeva, P. Tuberculosis Imaging, K. D. Kauffman, M. Donato, A. Sher, D. L. Barber, L. E. Via, T. J. Scriba, P. Khatri, Y. Song, K. W. Wong, C. M. Bosio, A. D. Klion, K. D. Mayer-Barber, Rapid GPR183-mediated recruitment of eosinophils to the lung after Mycobacterium tuberculosis infection. Cell Rep 40, 111144 (2022).

23. S. G. Hoft, M. A. Sallin, K. D. Kauffman, S. Sakai, V. V. Ganusov, D. L. Barber, The Rate of CD4 T Cell Entry into the Lungs during Mycobacterium tuberculosis Infection Is Determined by Partial and Opposing Effects of Multiple Chemokine Receptors. Infect Immun 87, (2019).

24. A. S. Hauser, M. M. Attwood, M. Rask-Andersen, H. B. Schioth, D. E. Gloriam, Trends in GPCR drug discovery: new agents, targets and indications. Nat Rev Drug Discov 16, 829–842 (2017).

25. S. H. Apte, M. E. Tan, V. P. Lutzky, T. A. De Silva, A. Fiene, J. Hundloe, D. Deller, C. Sullivan, P. T. Bell, D. C. Chambers, Alveolar crystal burden in stone workers with artificial stone silicosis. Respirology 27, 437–446 (2022).

26. Y. Hao, S. Hao, E. Andersen-Nissen, W. M. Mauck, 3rd, S. Zheng, A. Butler, M. J. Lee, A. J. Wilk, C. Darby, M. Zager, P. Hoffman, M. Stoeckius, E. Papalexi, E. P. Mimitou, J. Jain, A. Srivastava, T. Stuart, L. M. Fleming, B. Yeung, A. J. Rogers, J. M. McElrath, C. A. Blish, R. Gottardo, P. Smibert, R. Satija, Integrated analysis of multimodal single-cell data. Cell 184, 3573–3587 e3529 (2021).

27. G. Liu, M. A. Cooley, A. G. Jarnicki, T. Borghuis, P. M. Nair, G. Tjin, A. C. Hsu, T. J. Haw, M. Fricker, C. L. Harrison, B. Jones, N. G. Hansbro, P. A. Wark, J. C. Horvat, W. S. Argraves, B. G. Oliver, D. A. Knight, J. K. Burgess, P. M. Hansbro, Fibulin-1c regulates transforming growth factor-beta activation in pulmonary tissue fibrosis. JCI Insight 5, 124529–124546 (2019).

28. E. F. Redente, K. M. Jacobsen, J. J. Solomon, A. R. Lara, S. Faubel, R. C. Keith, P. M. Henson, G. P. Downey, D. W. Riches, Age and sex dimorphisms contribute to the severity of bleomycin-induced lung injury and fibrosis. Am J Physiol Lung Cell Mol Physiol 301, L510–518 (2011).

29. R. Lamichhane, S. Patial, Y. Saini, Higher susceptibility of males to bleomycin-induced pulmonary inflammation is associated with sex-specific transcriptomic differences in myeloid cells. Toxicol Appl Pharmacol 454, 116228–116258 (2022).

30. R. H. Hubner, W. Gitter, N. E. El Mokhtari, M. Mathiak, M. Both, H. Bolte, S. Freitag-Wolf, B. Bewig, Standardized quantification of pulmonary fibrosis in histological samples. Biotechniques 44, 507–511, 514-507 (2008).

