## Supplementary Materials for "Targeting the Oxysterol Receptor GPR183 to Mitigate Fibrogenesis in Idiopathic Pulmonary Fibrosis"

**Figure S1**

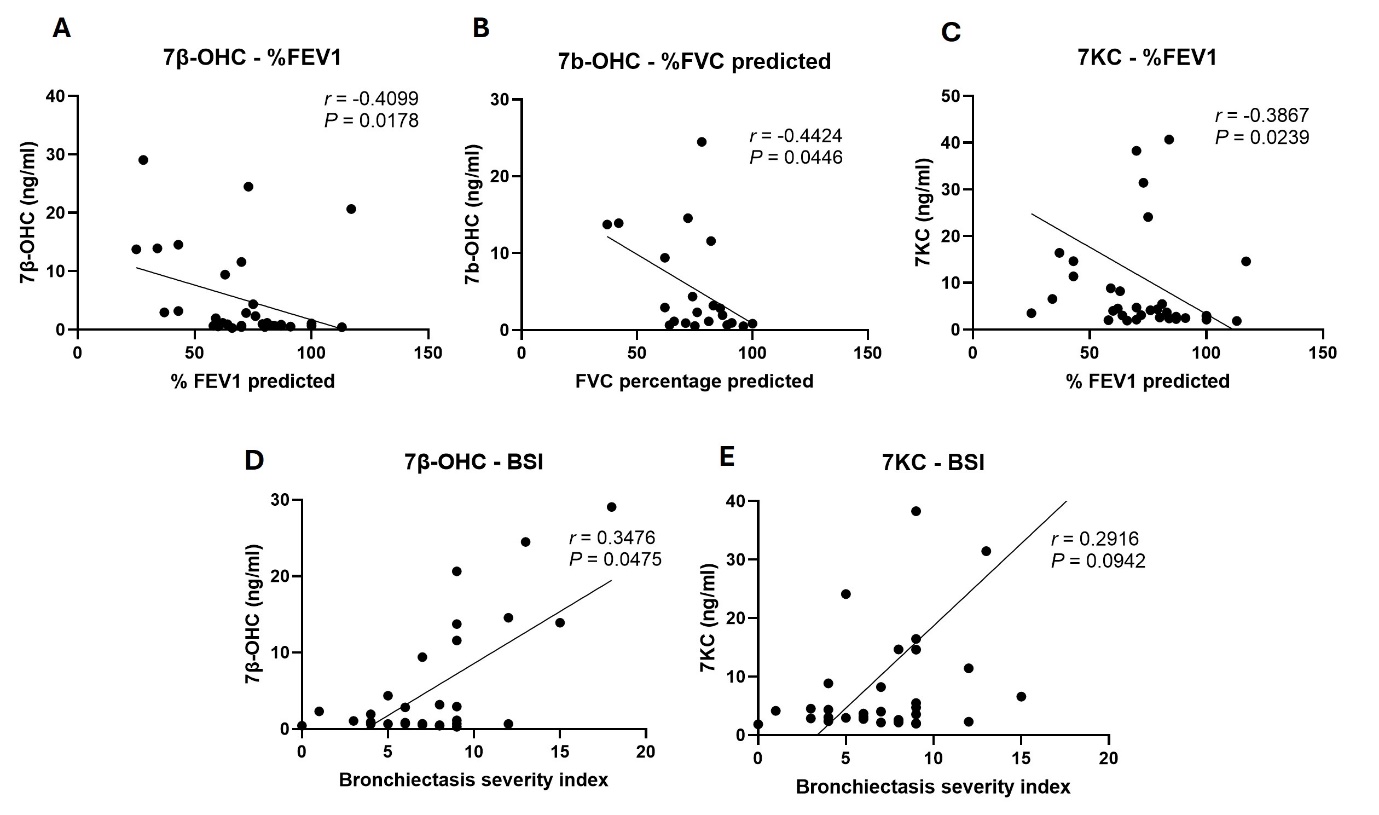
 **Fig. S1**. **Oxysterol concentrations are associated with lung function impairment and disease severity in bronchiectasis. (A-C)** Correlation analyses between BALF oxysterol concentrations and lung function parameters in patients with bronchiectasis. Relationships between 7β-hydroxycholesterol (7β-OHC) concentrations and percentage predicted FEV1 **(A)** and percentage FVC **(B)**, and between 7KC concentrations and percentage predicted FEV1 (C). **(D,E)** Correlation analyses between BALF oxysterol concentrations and BSI scores. Relationships between 7β-OHC concentrations and BSI **(D)** and between 7KC concentrations and BSI **(E)**. Spearman's rank correlation test was used to assess associations between variables (*n* = 34; clinical data were unavailable for 3 of 37 patients with bronchiectasis). Spearman correlation coefficients (*r*) and corresponding *P* values are shown in each panel. 7β-OHC, 7β-hydroxycholesterol; 7KC, 7-ketocholesterol; BALF, bronchoalveolar lavage fluid; FEV1, forced expiratory volume in 1 s; FVC, forced vital capacity; BSI, Bronchiectasis Severity Index.

**Figure S2**

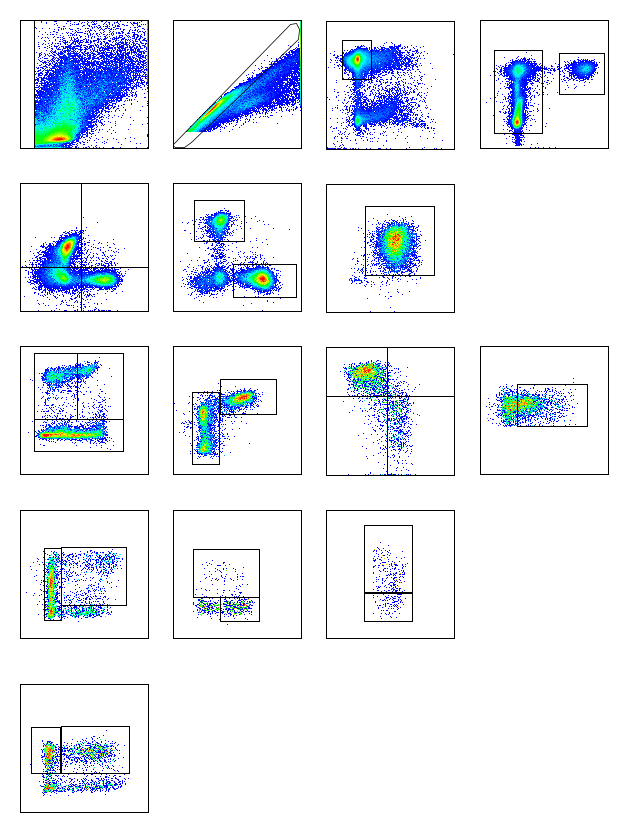

SSC-H

FSC-A

FSC-H

FSC-A

CD45

Viability

CD11b

Ly6G

**G1**

**G2**

**G3**

**G5**

**G6**

**G4**

**G5**

**G6**

**G4**

CD3

CD45R

CD4

CD8

CD24

MHCll

SigF

CD11c

**G8**

**G9**

**G7**

**G8**

CD11b

F4/80

Ly6C

CD11c

**G11**

F4/80

CD206

**G15**

**G13**

**G10**

CD11c

MHCll

CD103

CD11b

CD24

CD11b

**G12**

**G14**

**G14**

**Fig. S2. Flow cytometry gating strategy used to identify pulmonary immune cell populations.** Representative flow cytometry plots illustrating the gating strategy for immune cells in lung single cells suspension. Gates containing multiple cell populations are numbered (G1-G15). Immune cells are immunophenotyped as: Neutrophils (CD11b^+^ Ly6G^+^; G3), T cells (Ly6G^-^CD3^+^; G4), CD4^+^ T cells (G5), CD8^+^ T Cells (G5), B cells (Ly6G^-^CD45R^+^; G4), CD24^+^MHCII^+^ B cells (G6), Eosinophils (Ly6G^-^,B220^-^,CD3^-^,CD11c^-^,SigF^+^; G7), Alveolar macrophages (Ly6G^-^,B220^-^,CD3^-^,CD11c^+^,SigF^+^; G7), Macrophages (Ly6G^-^,B220^-^,CD3^-^,SigF^-^,CD11b^+^F4/80^+^, G8), CD11c^-^Ly6C^+^ macrophages (G9), CD11c^+^Ly6C^+^ macrophages (G9), CD11c^+^Ly6C^-^ macrophages (G9). CD206^+^ macrophages (G11), DCs (Ly6G^-^,B220^-^,CD3^-^,SigF^-^,CD11c^+^,MHCll+; G10), CD103 DCs (G13), CD24^+^cDC1 (CD103^-^,CD11b^+^,CD24^-^; G14), CD24^-^cDC2 (CD103^-^,CD11b^+^,CD24+; G14), Ly6C^-^ monocytes (Ly6G^-^,B220^-^,CD3^-^,SigF^-^, MHCll^-^,CD11b^+^,Ly6C^-^;G15) and Ly6C^+^ monocytes (Ly6G^-^,B220^-^,CD3^-^,SigF^-^, MHCll^-^,CD11b^+^,Ly6C^+^;G15).

**Figure S3**

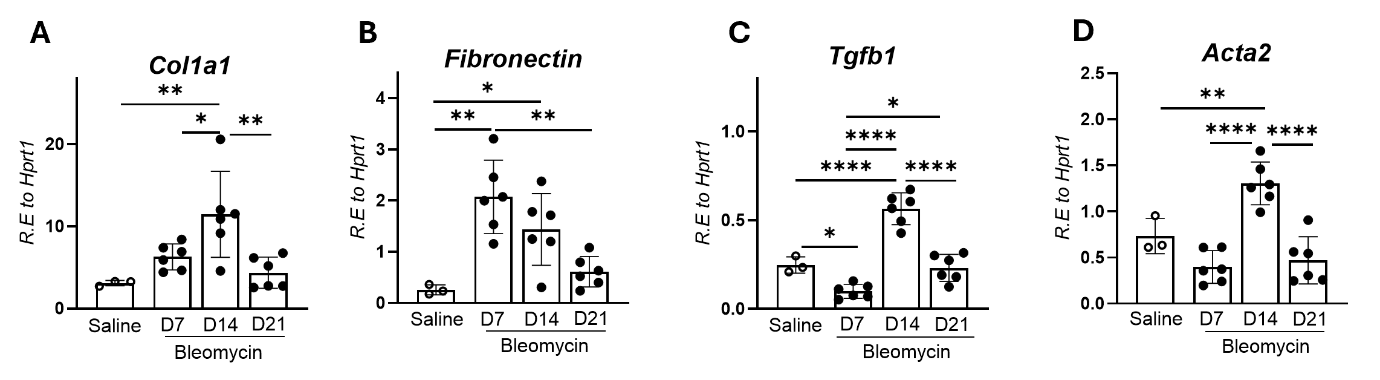
**Fig. S3. Expression of fibrosis-associated gene during bleomycin-induced pulmonary fibrosis. (A-D)** Relative lung mRNA expression of Col1a1 (**A**), Fn1 (**B**), Tgfb1 (**C**), and Acta2 (**D**) in whole-lung homogenates collected from saline-treated mice and mice at days 7, 14, and 21 following bleomycin administration. Gene expression was determined by quantitative PCR and normalized to Hprt1 expression. Data are presented as relative expression (R.E.) to Hprt1. Bars represent mean ± SD. Saline-treated mice (*n* = 3) and bleomycin-treated mice (*n* = 6 per time point). Statistical significance was assessed using ordinary one-way ANOVA followed by Tukey's multiple-comparison test. Exact statistical comparisons are indicated in the figure. **p* < 0.05, ***p* < 0.01, *****p* < 0.0001. R.E., relative expression; Col1a1, collagen type I alpha 1 chain; Fn1, fibronectin 1; Tgfb1, transforming growth factor-β1; Acta2, alpha-smooth muscle actin; Hprt1, hypoxanthine phosphoribosyltransferase 1.

**Figure S4**

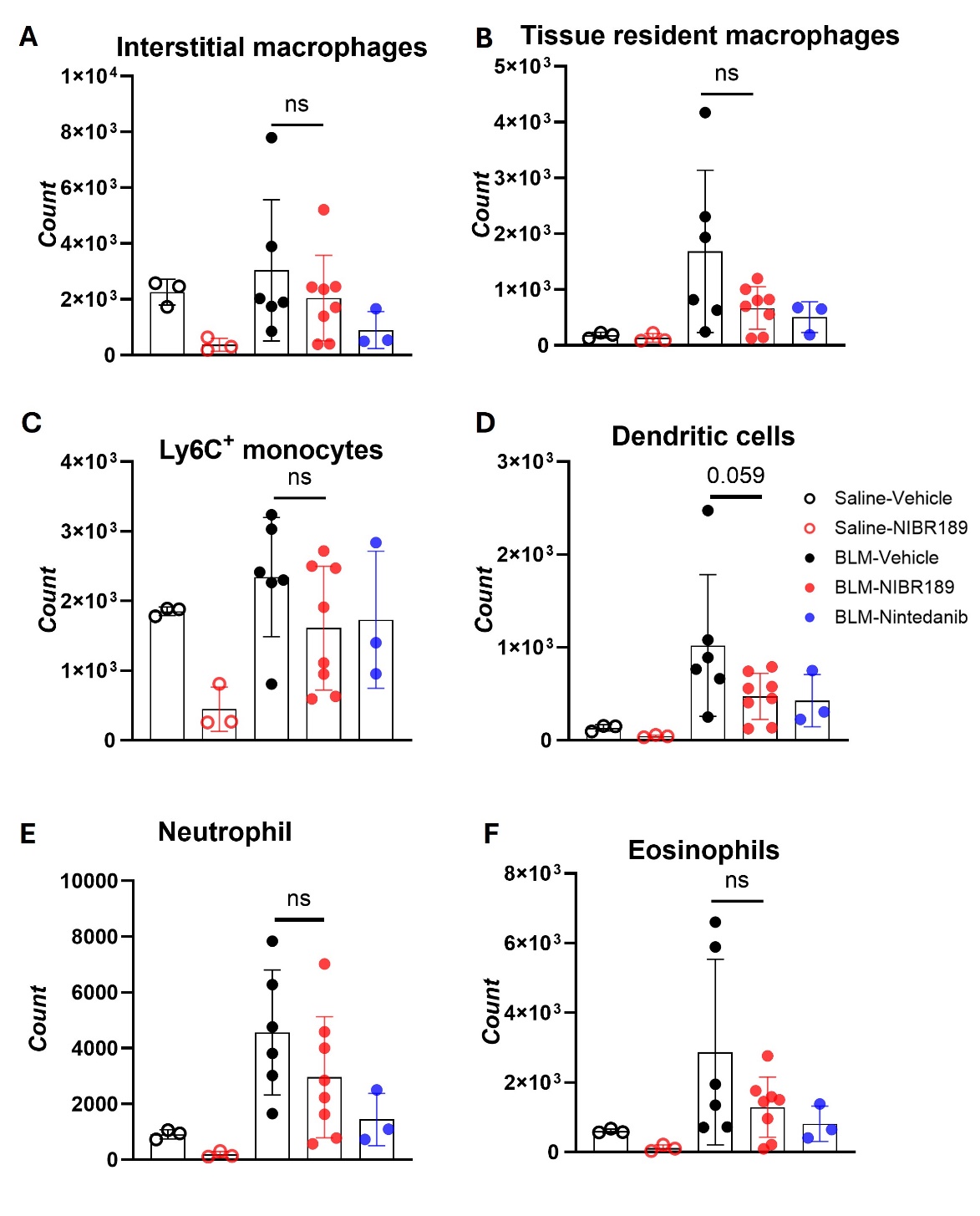

**Fig. S4. Effects of therapeutic GPR183 antagonism on major pulmonary myeloid cell populations following bleomycin-induced pulmonary fibrosis. (A-F)** Flow cytometric quantification of **(A)** interstitial macrophages, **(B)** tissue-resident alveolar macrophages, **(C)** Ly6C⁺ monocytes, **(D)** dendritic cells, **(E)** neutrophils, and **(F)** eosinophils in lungs collected at day 21 following BLM administration. Mice received vehicle (*n* = 6), the GPR183 antagonist NIBR189 (*n* = 8), or nintedanib (*n* = 3) beginning on day 10 after BLM administration. Saline-treated mice served as controls (n = 3 per treatment). Bars represent mean ± SD. Statistical significance was assessed using Kruskal-Wallis analysis followed by Dunn’s multiple-comparison test. ns, not significant.

**Figure S5**

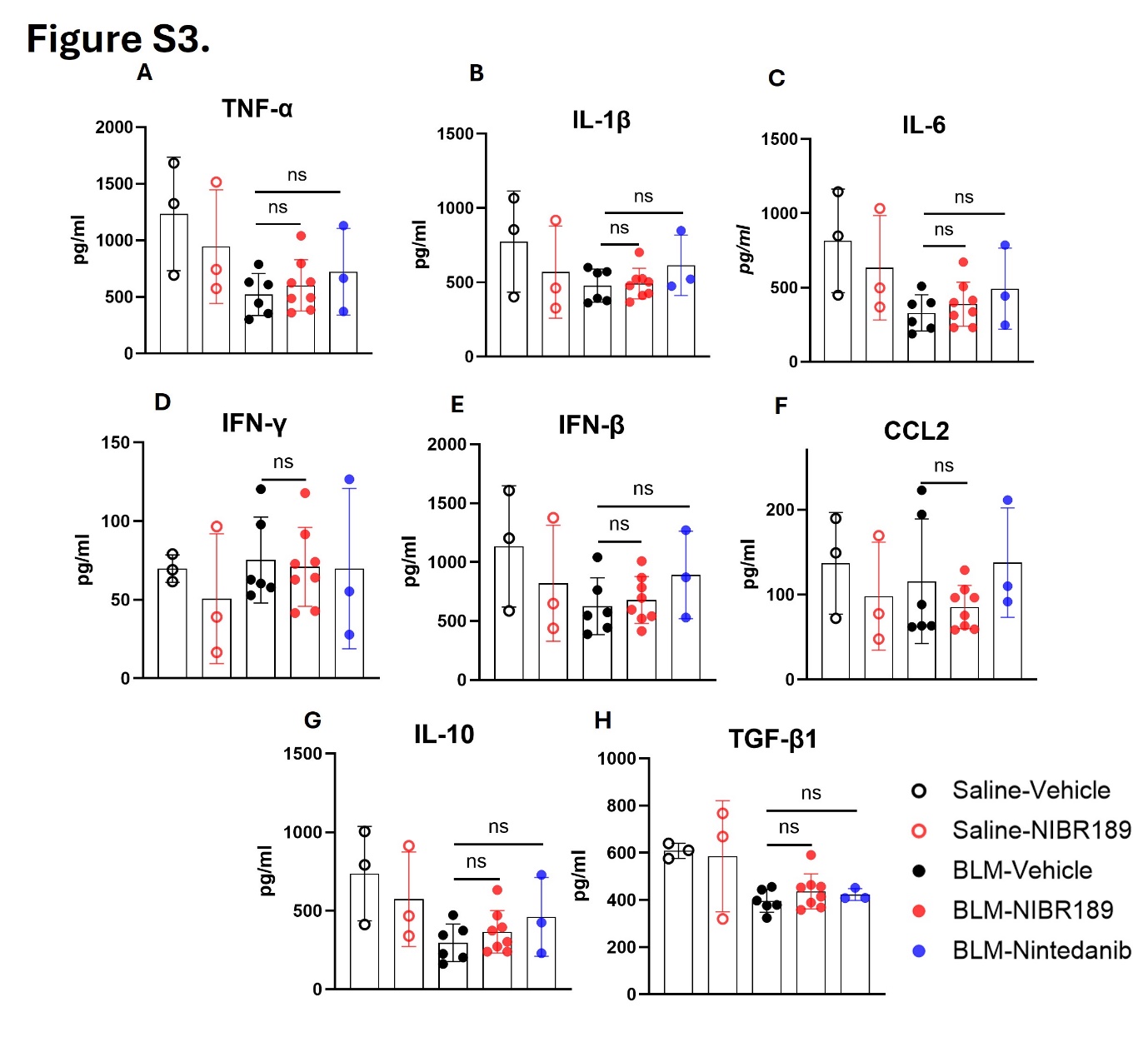

**Fig. S5. Concentrations of lung cytokines post therapeutic GPR183 antagonism has limited effects on** **pulmonary cytokine concentrations during bleomycin-induced pulmonary fibrosis.**

Lung concentrations of tumour necrosis factor-α (TNF-α; **A**), interleukin-1β (IL-1β; **B**), interleukin-6 (IL-6; **C**), interferon-γ (IFN-γ; **D**), interferon-β (IFN-β; **E**), C-C motif chemokine ligand 2 (CCL2; **F**), interleukin-10 (IL-10; **G**), and transforming growth factor-β1 (TGF-β1; **H**) measured by ELISA in lung homogenates collected at day 21 following BLM administration. Mice received saline or BLM and were treated with vehicle, the GPR183 antagonist NIBR189, or nintedanib as indicated. Open symbols represent saline-treated mice and filled symbols represent BLM-treated mice. Black symbols indicate vehicle-treated mice, red symbols indicate NIBR189-treated mice, and blue symbols indicate nintedanib-treated mice. Bars indicate mean ± SD. Statistical significance was determined using Kruskal-Wallis analysis followed by Dunn's multiple-comparison test. Exact statistical comparisons are shown in the figure. *ns*, not significant.

**Figure S6**

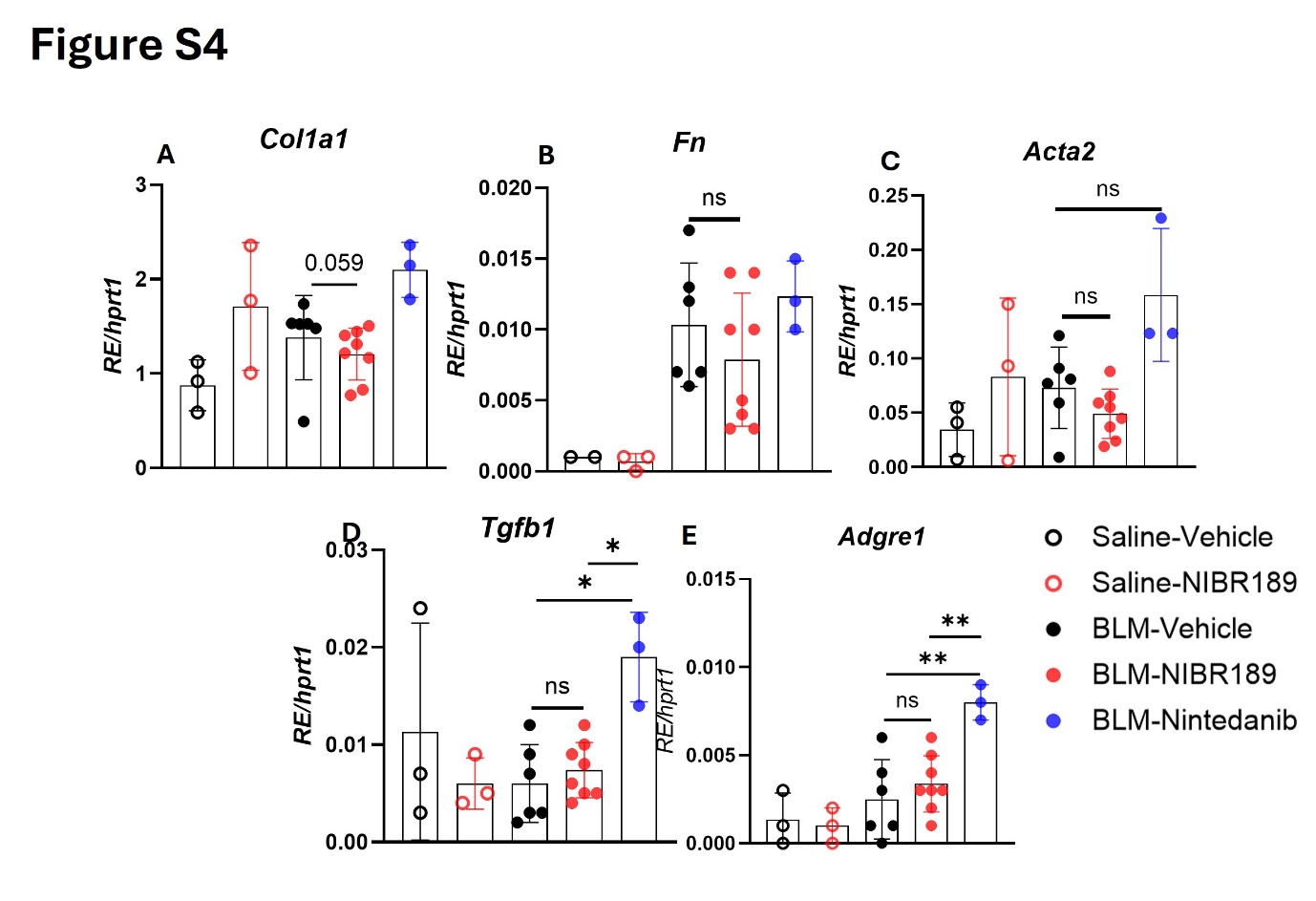

**Fig. S6. Therapeutic GPR183 antagonism modulates expression of fibrosis- and macrophage-associated genes during bleomycin-induced pulmonary fibrosis.**

Relative lung mRNA expression of the fibrosis-associated genes Col1a1 (**A**), Fn1 (**B**), Acta2 (**C**), Tgfb1 (**D**), and the macrophage marker Adgre1 (**E**) in saline-treated and bleomycin (BLM)-treated mice receiving vehicle, NIBR189, or nintedanib. Gene expression was determined by quantitative PCR and normalized to Hprt1 expression. Bars indicate mean ± SD. Statistical significance was determined using Kruskal-Wallis analysis followed by Dunn’s multiple-comparison test. Exact statistical comparisons are shown in the figure. **p* < 0.05, **p < 0.01; ns, not significant.

**Supplement table 1.** Optimized MRM transitions and oxysterol-specific mass spectrometry parameters used in the data acquisition method for quantification.

| **Analyte** | **Q1 Mass (Da)** | **Q3 Mass (Da)** | **RT (min)** | **DP (V)** | **EP (V)** | **CE (V)** | **CXP (V)** |
| --- | --- | --- | --- | --- | --- | --- | --- |
| 7α,25-OHC | 383.20 | 365.30 | 3 | 120 | 10 | 30 | 7 |
| 7α,25-OHC | 383.20 | 147.30 | 3 | 120 | 8 | 30 | 7 |
| 7α,25-OHC | 383.20 | 159.00 | 3 | 120 | 8 | 30 | 7 |
| 25-OHC | 385.30 | 367.33 | 6.4 | 80 | 10 | 20 | 14 |
| 24-OHC | 385.30 | 367.33 | 6.8 | 80 | 10 | 20 | 14 |
| 27-OHC | 385.30 | 147.10 | 7 | 80 | 10 | 50 | 15 |
| 27-OHC | 385.30 | 367.33 | 7 | 80 | 10 | 20 | 14 |
| 7β-OHC | 385.31 | 367.30 | 8.9 | 70 | 7 | 25 | 15 |
| 7-ketocholesterol | 401.30 | 383.40 | 10.4 | 90 | 7 | 30 | 15 |

**Abbreviations:** RT, retention time; DP, declustering potential; EP, entrance potential; CE, collision energy; CXP, collision cell exit potential.

**Supplement table 2. Antibodies used for Flow cytometry**

| **Anti-mouse antibodies** | **Identifier** |
| --- | --- |
| VioletFluor 450 – CD11b | 75-0112-U100 |
| PE-Cy7 – CD11c | 60-0114-U100 |
| PerCP-Cy5.5 – Ly6G | 65-1276-U100 |
| VioletFluor 500 – MHCII | 85-5321-U100 |
| PerCP – CD4 | 67-0042-U100 |
| RedFluor 710 – CD45R | 80-0452-U100 |
| ViaDye Red - Viability | R7-60008 |
| BUV737 – CD103 | 367-1031-82 |
| BUV805 – CD24 | 368-0242-82 |
| Super Bright 702 – CD3 | 67-0032-82 |
| BUV496 – CD8 | 364-0081-82 |
| Alexa Fluor 488 – F4/80 | 53-4801-82 |
| PE – CD163 | 12-1631-82 |
| Alexa Fluor 647 – SigF | 155520 |
| APC-Fire 750 – CD14 | 123332 |
| PE/Dazzle – Ly6C | 128044 |
| BUV395 – CD45 | 564279 |
| PE – CD206 | 568273 |

**Supplement table 3.** Primer sequences for qRT-PCR

| **Gene** | **Forward primer** | **Reverse primer** |
| --- | --- | --- |
| *Col1a1* | CTTCACCTACAGCACCCTTGTG | TGACTGTCTTGCCCCAAGTTC |
| *Fn* | TGTGGTTGCCTTGCACGAT | GCTATCCACTGGGCAGTAAAGC |
| *Acta2* | CCCAGACATCAGGGAGTAATGG | TCTATCGGATACTTCAGCGTCA |
| *Tgfb1* | CCCGAAGCGGACTACTATGCTA | GGTAACGCCAGGAATTGTTGCTAT |
| *Gpr183* | GTCGTGTTCATCCTGTGCTTCAC | TCATCAGGCACACCGTGAAGTG |
| *Ch25h* | CTGACCTTCTTCGACGTGCT | GGGAAGTCATAGCCCGAGTG |
| *Cyp7b1* | CGGAAATCTTCGATGCTCCAAAG | GCTTGTTCCGAGTCCAAAAGGC |
| *Adgre1* | CGTGTTGTTGGTGGCACTGTGA | CCACATCAGTGTTCCAGGAGAC |
| *Hprt1* | CTGGTGAAAAGGACCTCTCGAAG | CCAGTTTCACTAATGACACAAACG |
